# Microglia Detect Externalized Phosphatidylserine on Synapses for Elimination via TREM2 in Alzheimer’s Disease Models

**DOI:** 10.1101/2022.04.04.486424

**Authors:** Javier Rueda-Carrasco, Dimitra Sokolova, Sang-Eun Lee, Thomas Childs, Natália Jurčáková, Sebastiaan De Schepper, Judy Z. Ge, Joanne I. Lachica, Christina E. Toomey, Oliver J. Freeman, John Hardy, Beth Stevens, Tammaryn Lashley, Sunghoe Chang, Soyon Hong

**Author notes:** These authors contributed equally to this work (Javier Rueda-Carrasco, Dimitra Sokolova, Sang-Eun Lee).

## Abstract

Genetic studies implicate phagocytosis pathways in microglia to be a major Alzheimer’s disease (AD)-associated process. Microglia phagocytose synapses in AD mouse models, suggesting a role for microglia in region-specific synapse loss, a pathological hallmark of AD. However, whether specific synapses are targeted for elimination, and if so, how, remains to be elucidated. Here, we show that synapses externalize phosphatidylserine (PtdSer) upon challenge by β-amyloid oligomers, which are then selectively engulfed by microglia. Mechanistically, we find that Triggering Receptor Expressed on Myeloid Cells 2 (TREM2) is critical for microglia to sense and preferentially engulf AD synapses. In brains of mice and humans, TREM2 dysfunction leads to exacerbation of apoptotic synapses. Our work altogether suggests a fundamental role for microglia as brain-resident macrophages to remove damaged synapses in AD. We provide mechanistic insight into how TREM2 variants associated with increased risk of developing AD may contribute to defective microglia-synapse function.

**One-Sentence summary:** Microglia selectively engulf synapses in Alzheimer-like mouse brains via PtdSer-TREM2 signaling.

## Main Text

Region-specific loss and dysfunction of synapses is an early pathological hallmark of AD and a major correlate of cognitive impairment (*1*). Genetic studies in sporadic AD implicate microglia, the primary tissue-resident macrophages of the brain, as significant modulators for AD risk (*2–4*). However, how microglia contribute to AD risk remains elusive. Microglia are active participants in brain wiring, where they sculpt neural circuits and influence synaptic refinement and function (*5*, *6*). In various models of disease, including those of AD, microglia-synapse interactions are dysregulated, leading to synaptic loss (*7*). However, whether specific synapses are vulnerable to microglial engulfment in AD is unknown.

A fundamental role of tissue-resident macrophages is to selectively remove unwanted materials in their local microenvironment for proper maintenance of tissue homeostasis (*8*). Phosphatidylserine (PtdSer), a phospholipid enriched on cell membranes, is a highly conserved ‘eat me’ signal expressed on apoptotic cells. Normally asymmetrically localized to the inner leaflet of plasma membranes, PtdSer is externalized (ePtdSer) on the outer surface of membranes during caspase-mediated apoptosis, marking the cell for selective removal by phagocytes (*9*). Interestingly, in the developing brain during peak synapse pruning period, synapses locally express ePtdSer for engulfment by microglia (*10, 11*). Similar focal apoptotic-like mechanisms on synapses, in a process known as synaptosis, have been proposed in mouse models of AD (*12–14*). However, whether ePtdSer is upregulated on synapses in AD to govern selective microglial engulfment is not known. Here, we hypothesized that synaptic ePtdSer underlies vulnerability of synapses to microglial engulfment in AD.

To address whether synaptic ePtdSer is increased in AD-relevant context and acts as ‘eat me’ signals for microglia, we first visualized PtdSer externalization on Homer1-eGFP (*15*) synaptic membranes in real-time following exposure to β-amyloid oligomers (oAβ), which have been shown to impair synapses (*16–18*) and stimulate microglia-synapse engulfment *in vivo* (*19*). Application of a physiologically relevant concentration (50 nM) of soluble oAβ to primary mouse hippocampal neurons resulted in rapidly elevated levels of ePtdSer on Homer1-eGFP^+^ dendritic spines (**Fig. 1A** and **B**), as shown by PSVue bis-Zinc^2+^-dipicolylamine (Zn-DPA) which binds with high affinity to ePtdSer (*10*, *20*). There were negligible levels of PSVue signal on eGFP^+^ spines in control conditions during this time period, suggesting healthy basal culture levels (**Fig. S1A**). Further, PSVue did not fluoresce across the whole neuron, but was found to align with Homer1-eGFP (**Fig. S1A** and **B**). Super-resolution microscopy (SRM) imaging confirmed punctate PSVue colocalization with Homer1-eGFP^+^ (**Fig. 1C**) and along oAβ-bound PSD95-immunostained dendrites (**Fig. S1C**), suggesting that oAβ’s deposition on synapses (*17*) induces focal ePtdSer. Indeed, we observed a markedly higher percentage of Homer1-eGFP to be colocalized with PSVue upon oAβ treatment (**Fig. 1D**); likewise, a higher proportion of PSVue was found to colocalize with Homer1-eGFP (**Fig. 1E**). Altogether, these results suggest that synapses externalize PtdSer upon oAβ challenge.

**Fig. 1.**
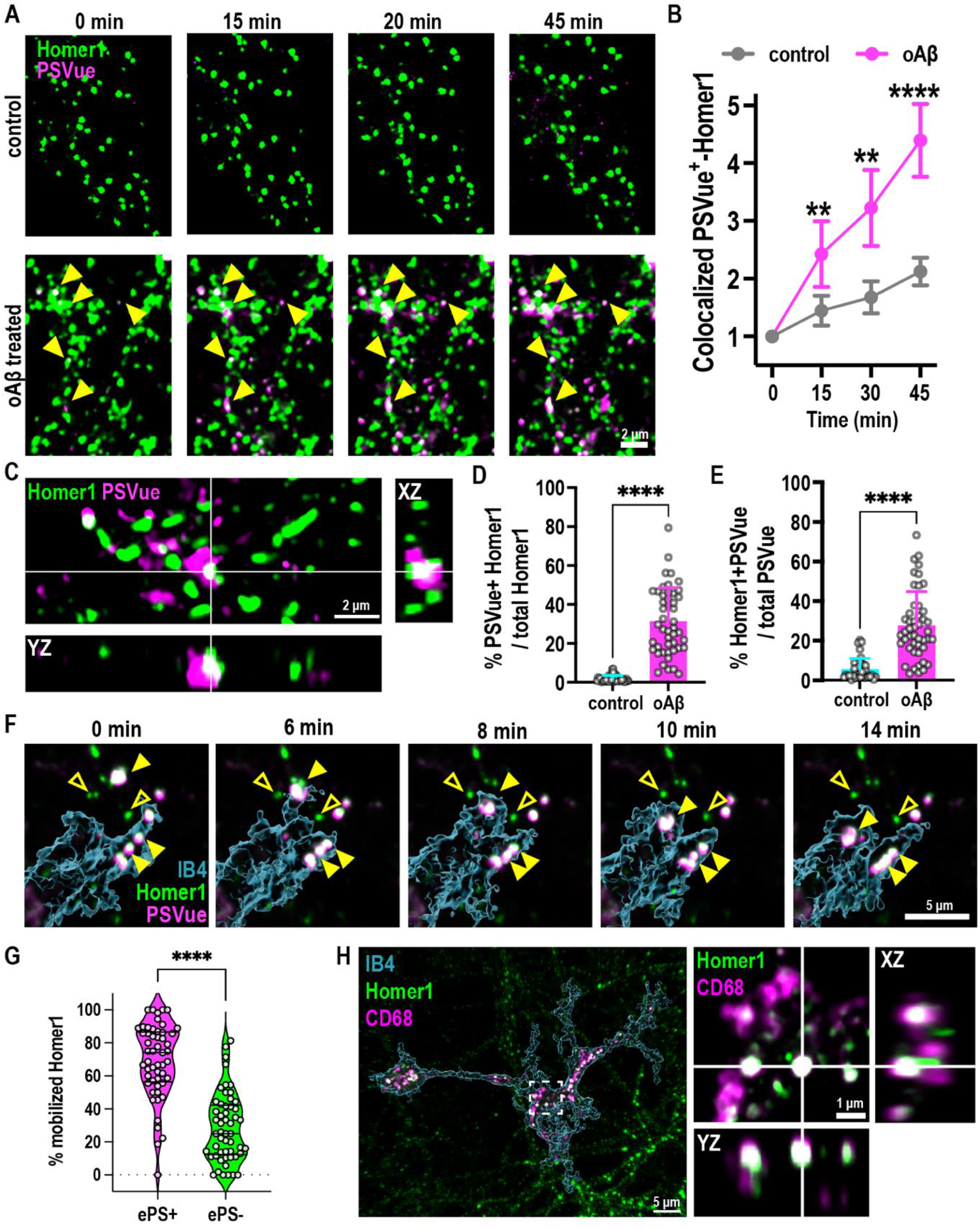
Microglia target ePtdSer^+^ spines for engulfment upon oAβ challenge. **A)** Time-lapse images of primary Homer1-eGFP neurons (green) treated with 50 nM oAβ versus vehicle control. Yellow arrows indicate increasing PSVue550 (magenta) signal on dendritic spines with oAβ treatment over 45 min. **B)** Relative fold-change of colocalized PSVue and Homer1-eGFP signal with time of **(A)**. **C)** SRM images of Homer1-eGFP neurons and PSVue at 1 h post oAβ treatment. **D-E**) Quantification of colocalized PSVue and Homer1 -eGFP shown as percentage of total Homer1-eGFP (**D**) or total PSVue (**E**). **F)** Time-lapse images of microglia (cyan, labelled with IB4-647, 3D rendered) internalizing (yellow arrowheads) PSVue^+^ Homer1-eGFP dendritic spines. Note PSVue^-^ Homer1-eGFP dendritic spines are left behind (empty yellow arrowheads). **G)** Index of Homer1 puncta in motion within 10 min of recording show that microglia preferentially pull Homer1-eGFP^+^ PSVue^+^ (75%) over Homer1-eGFP^+^ PSVue^-^ (25%). **H**) SRM images of Homer1-eGFP and CD68 (magenta) lysosomes in microglia. Inset shows orthogonal view of colocalized Homer1-eGFP and CD68. Data shown as mean ± SD, 1 point represents 1 ROI. n=24 **(A-B)**, 38 (control), 47 (oAβ) **(D-E)** and 30 **(G)** ROIs from 3-5 independent culture sets. Two-way ANOVA followed by Tukey post-hoc test **(A)**, unpaired t-test **(D, E** and **G)**, p-values **P<0.01; ****P<0.0001.

To address whether microglia selectively target ePtdSer^+^ spines for phagocytosis, we established a microglia-neuron co-culture using hippocampal neurons from Homer1-eGFP mice allowing us to image microglia-neuron interactions at the level of spines in real-time. Primary microglia were supplemented with mCSF1, TGF-β1 and CX3CL1 in order to induce a more homeostatic microglial transcriptome in culture (*21*, *22*) (**Fig. S2A**). Using live-cell imaging with 3D high-resolution capacity, we found that microglial processes primarily contact (**Fig. S2B** and **movie S1**) and engulf (**Fig. 1F** and **movie S2**) the Homer1-eGFP^+^ spines that become PSVue^+^ upon oAβ challenge. Interestingly, microglial processes appear to target and contact PSVue^+^ eGFP^+^ spines for some time (**Fig. S2B**) whereas engulfment itself appears to be a more rapid event (**Fig. 1F**). Importantly, microglia preferentially engulfed PSVue^+^ eGFP^+^ over PSVue^-^ eGFP^+^ synaptic elements (**Movie S2** and **fig. 1G**), suggesting selectivity for the engulfment process. SRM showed engulfed Homer1-eGFP^+^ inside CD68^+^ lysosomes in microglia (**Fig. 1H**). Altogether, these data suggest that microglia target ePtdSer^+^ synapses for removal.

We thus directly tested whether there is selectivity to microglia-synapse engulfment and the role of synaptic ePtdSer in this process. We first asked whether certain synapses are more vulnerable to microglial engulfment. Treatment of freshly prepared wild-type (WT) mouse brain synaptosomes (*23*) with oAβ resulted in an increase of ePtdSer on outer membranes (**Fig. S3A**, **B**, **C** and **D**). Microglia rapidly and preferentially engulfed oAβ-treated synaptosomes over control synaptosomes (**Fig. 2A, B** and **C**); this discrimination by microglia lasted over 10 h of live-cell time-lapse imaging (**Movie S3**). We confirmed that the microglial preference for oAβ-treated synaptosomes was independent of pHrodo dye conjugates used (**Fig. S4A**, **B** and **C**) and that fluorescence signals were pH-dependent (**Fig. S4D** and **E**). Similarly, microglia preferentially engulfed synaptosomes isolated from frontal cortex of patient brains with AD pathology versus those of non-demented controls (NDC) (**Fig. 2D** and **table S2**); AD synaptosomes contained a significantly higher level of Aβ monomers and dimers versus synaptosomes from NDC (**Fig. S3E**). To test whether ePtdSer signals are required for microglial engulfment, we incubated oAβ-treated mouse and human AD synaptosomes with Annexin-V (AnnxV), which masks ePtdSer (*24*). There was a marked decrease of engulfment of both oAβ-treated mouse and human AD synaptosomes by microglia (**Fig. 2D, E** and **fig. S5A**, **B** and **C**). Altogether, these data suggest that microglia preferentially target AD synapses for engulfment, and that the specificity is promoted by ePtdSer on synaptosomes.

**Fig. 2.**
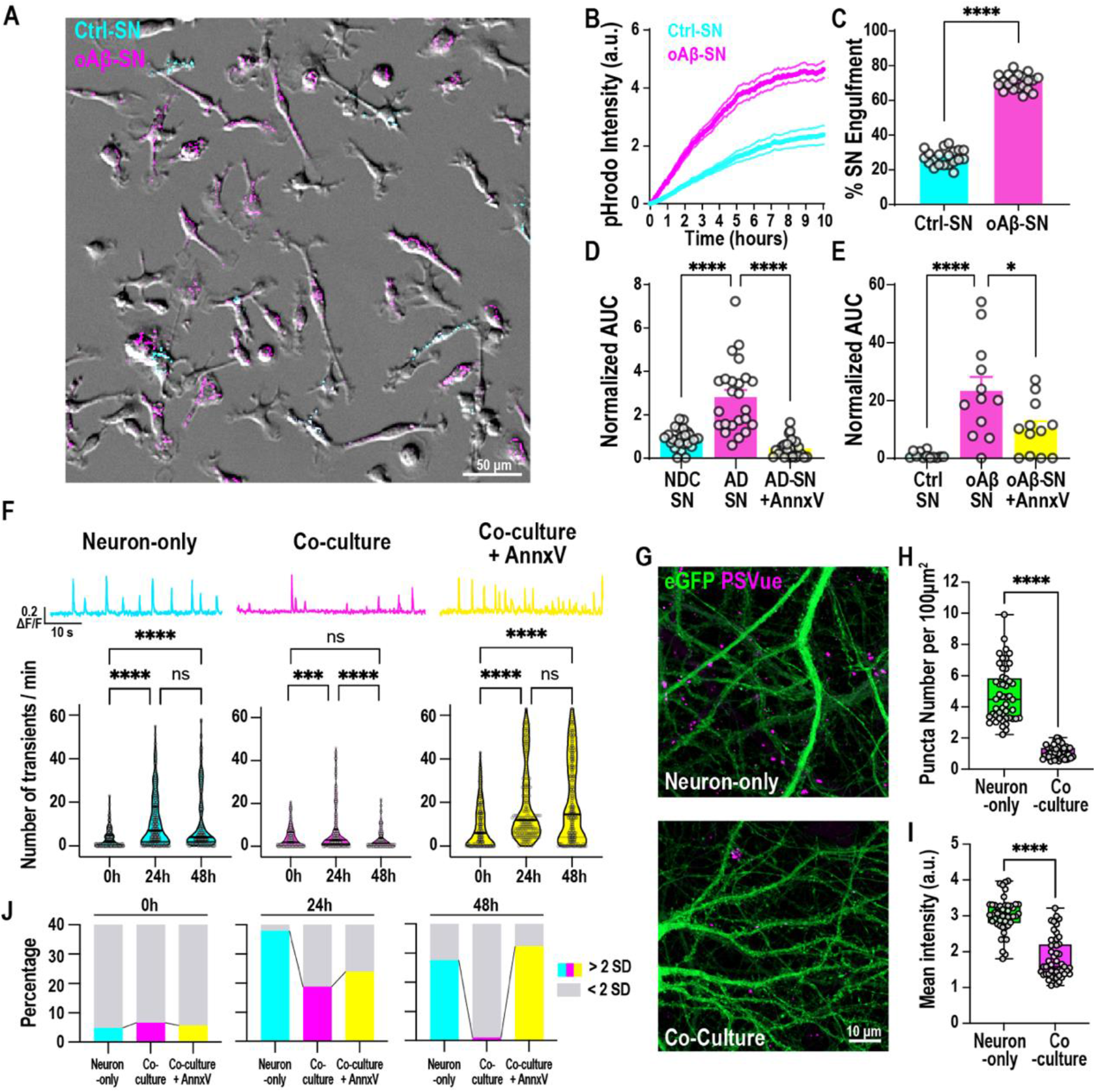
Microglia engulf mouse and human AD synaptosomes via ePtdSer. **A)** Primary microglia simultaneously treated with oAβ-synaptosomes (oAß-SN) in pHrodo red (magenta) and control synaptosomes (Ctrl-SN) in pHrodo deep red (cyan). **B)** pHrodo fluorescence (a.u.) over 10 h (3-5 min intervals, synaptosomes added at t=0) showing a faster rate of increase of oAβ-SN compared to Ctrl-SN. **C)** Percent of total engulfed pHrodo fluorescence that reflects oAβ-SN or Ctrl-SN at t=3 h. **D-E)** pHrodo fluorescence with time shown as area under curve (AUC) at 3 h normalized to respective control. AUC of engulfed human AD SN versus NDC SN **(D)** and mouse oAβ-SN versus Ctrl-SN **(E)**, with and without AnnxV pre-treatment. N=6 NDC versus AD patient cases. **F)** Representative trace (top panel) of transfected post-synaptic jGCaMP7 signal at 48 h post oAβ challenge and number of spontaneous calcium transients (bottom panel) per minute of neuron-only (cyan), neuron-microglia co-culture (magenta) or neuron-microglia co-culture treated with AnnxV (yellow) before (0 h) and after (24 h, 48 h). **G)** SRM images of neuron-only and neuron-microglia co-culture labelled with PSVue (magenta) 48 h after 50 nM oAβ treatment; neurons are transfected with jGCamp7c-eGFP (green). **H-I)** PSVue puncta per 100 μm^2^ **(H)** and mean intensity of PSVue **(I)** of neuron-only and neuron-microglia co-culture. **J)** Percentage of high frequency (cyan, magenta and yellow) and low frequency spines (grey) in each condition. High versus low, as defined by mean + 2 SD of the initial value. Data shown as mean ± SEM **(A-E)**. 1 point represents 1 ROI. 4-6 (40 cells per ROI) **(A-E)**, 60-130 **(F),** 16 **(I-J)** ROIs per experiment, n=3 experiments. Paired t-test **(C)**, one-way ANOVA followed by Bonferroni post-hoc test **(D, E)** or Kruskal-Wallis test followed by Dunn’s multiple comparisons test **(F)**, Unpaired t-test **(H-I)**, p-values shown ns P>0.05; *P<0.05; **P<0.01; ***P<0.001; ****P<0.0001. Scale bar 50 μm **(A)** and 10 μm **(G)**.

We measured spontaneous calcium transients on dendrites of primary neuronal cultures (*25*) (**Movie S4**) and found that oAβ induced neuronal hyperactivity within 24 h. Interestingly, when neurons were co-cultured with microglia, neuronal hyperactivity decreased back to baseline levels within 48 h (**Fig. 2F** and **fig. S6**). Levels of prolonged hyperactivity corresponded to levels of PSVue signals left on dendrites: in microglia-neuron co-cultures, PSVue signals on dendrites decreased significantly as compared to those in neuron-only cultures (**Fig. 2G, H** and **I**). Importantly, the oAβ-induced hyperactivity continued to stay elevated when microglia-neuron co-cultures were treated with AnnxV (**Fig. 2F** and **fig. S6**), suggesting that ePtdSer is required for the resolution of neuronal hyperactivity by microglia. In-depth analysis showed that the fraction of spines that displayed high-frequency calcium transients (with frequency levels > mean + 2 SD of the initial value) were almost absent in microglia-neuron co-culture despite oAβ challenge, as compared to neuron-only cultures and to microglia-neuron co-culture treated with AnnxV (**Fig. 2J**). These results altogether suggest that microglia help regulate activity levels and raise the question of how microglia detect synaptic ePtdSer for phagocytosis.

TREM2, which is expressed on macrophages, has been shown to sense membrane damage-associated lipids including ePtdSer (*26*). TREM2 signaling induces transcriptional changes in macrophages towards highly phagocytic and lipid regulatory states to promote tissue homeostasis (*27*, *28*). Interestingly, TREM2 has been shown to mediate microglia-synapse engulfment during normal developmental synapse pruning (*29*) as well as in models of tau-mediated neurodegeneration (*30*). We thus asked whether the preferential engulfment for ePtdSer^+^ AD synapses by microglia is mediated via TREM2. To that end, we isolated microglia from Trem2 R47H knock-in (KI) mice where TREM2 expression (*31*) and function (*32*) are impaired. No gross differences in microglial motility or morphology were observed between WT and Trem2 R47H KI microglia (**Movie S5**). However, unlike WT microglia, Trem2 R47H KI microglia failed to engulf oAβ-treated synaptosomes (**Fig. 3A** and **B**), suggesting that functional TREM2 is required for the selective removal of ePtdSer^+^ Aβ^+^ synaptosomes.

**Fig. 3.**
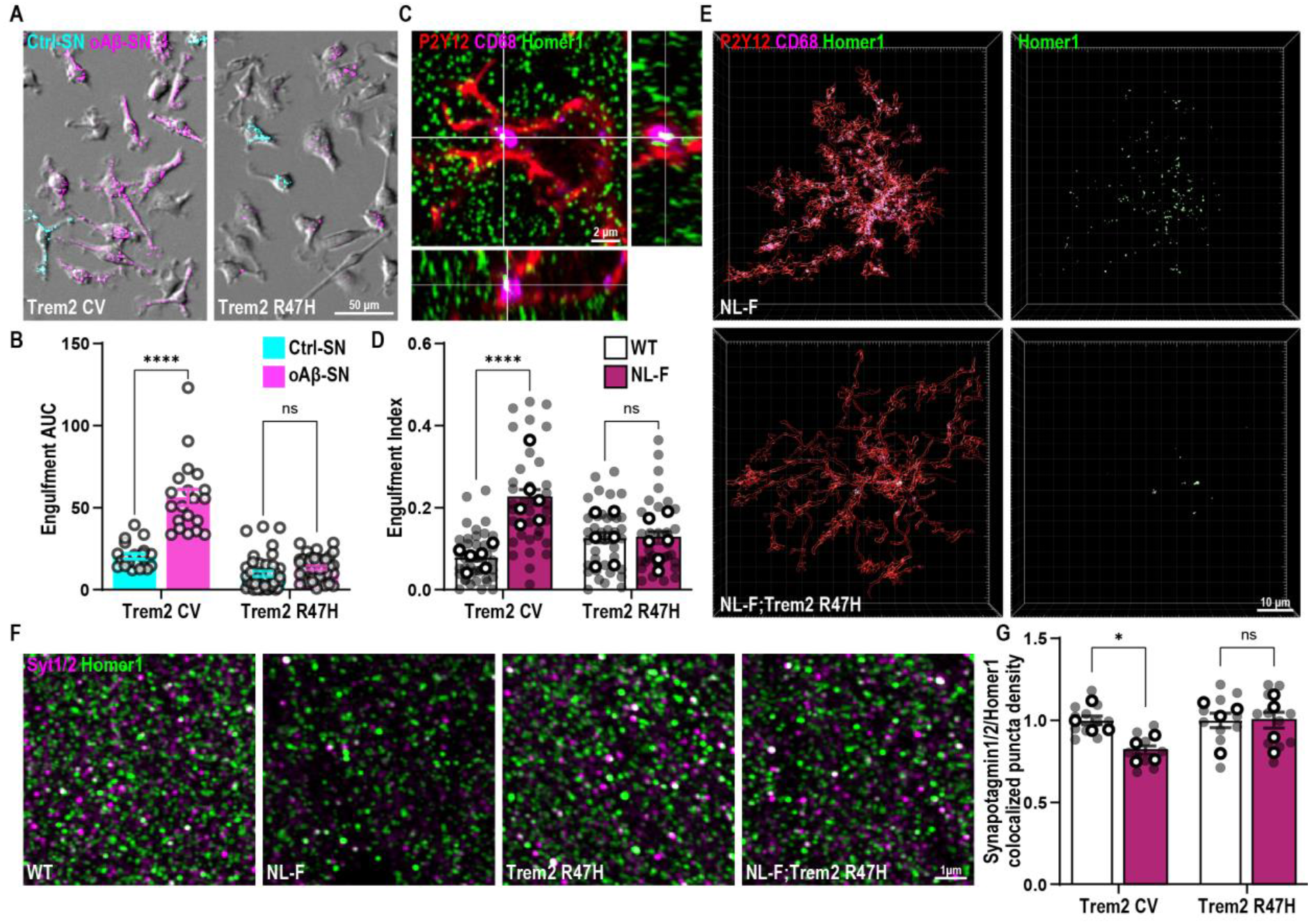
Loss-of-function TREM2 impairs microglial engulfment of synapses in AD models. **A)** Trem2 CV and Trem2 R47H KI primary microglia treated simultaneously with oAβ-synaptosomes (oAβ-SN) in pHrodo red (magenta) and control synaptosomes (Ctrl-SN) in pHrodo deep red (cyan). **B)** pHrodo fluorescence with time shown as area under curve (AUC) at 3 h. AUC of oAβ-SN is higher compared to Ctrl-SN in Trem2 CV but not in Trem2 R47H KI microglia. **C-E)** Hippocampal CA1 SR of 6 mo WT, Trem2 R47H KI, NL-F KI and NL-F KI; Trem2 R47H KI mice immunostained for P2Y12 (red), CD68 (magenta) and Homer1 (green). **C)** Orthogonal view showing engulfed Homer1 inside CD68 lysosomes in P2Y12 microglia. **D)** Quantification of engulfment index (volume of Homer1 in CD68/microglial cell volume)*100). **E)** Representative 3D surface rendering reconstructions showing increased Homer1 in microglia in NL-F KI compared to WT, Trem2 R47H KI, and NL-F KI; Trem2 R47H KI mice. **F)** SRM images from the hippocampal CA1 SR of 6 mo WT, Trem2 R47H KI, NL-F KI and NL-F KI; Trem2 R47H KI mice immunostained for Synaptotagmin (Syt1/2, magenta) and Homer1 (green), pre- and post-synaptic puncta, respectively. **G)** Colocalized puncta density normalized to WT or Trem2 R47H KI accordingly showing decreased synapse density in NL-F KI but not NL-F KI; Trem2 R47H KI. Data shown as mean ± SEM. 1 point represents 1 ROI (40 cells per ROI), 6-12 ROIs per experiment, n=3 experiments **(B)**. Shaded points represent 1 ROI and open points represent animal average **(D**, **G)**. 6-9 cells (1 ROI represents 1 cell) per animal **(D)**, 3 ROIs per animal **(G)**, n=4-6 animals. Two-way ANOVA followed by Bonferroni’s post-hoc test, p-values shown as ns P>0.05; *P<0.05; ****P<0.0001. Scale bar 50 μm **(A)**, 2 μm **(C)**, 10 μm **(E)** and 1 μm **(E).**

To address *in vivo* relevance, we crossed Trem2 R47H KI line with the slow progressing hAPP NL-F KI mice, which allows for endogenous control of Aβ production (*33*). Using *in vivo* microglia-synapse engulfment assay (*19*), we found in 6 mo NL-F mice, an age that precedes robust plaque deposition (*33*), significantly higher volumes of engulfed Homer1-immunoreactive synaptic puncta inside CD68^+^ lysosomes of P2Y12^+^ microglia in the hippocampal CA1 strata radiatum (SR) as compared to those in age- and sex-matched WT controls (**Fig. 3C, D, E** and **fig. S7)**. SRM imaging also showed loss of colocalized Homer1- and Synaptotagmin1/2-immunoreactive synaptic puncta in the hippocampal CA1 SR in the 6 mo NL-F hippocampus versus WT controls (**Fig. 3F** and **G**).

Conversely, in 6 mo NL-F;Trem2 R47H KI mice, levels of engulfed Homer1^+^ synaptic puncta by microglia were comparable to those of age- and sex-matched Trem2 R47H KI mice, suggesting that microglia with dysfunctional TREM2 fail to engulf synapses *in vivo* as well as *in vitro,* despite Aβ challenge (**Fig. 3**). Levels of Homer1- and Synaptotagmin1/2-immunoreactive synaptic puncta were comparable between NL-F;Trem2 R47H KI and Trem2 R47H KI hippocampus (**Fig. 3F** and **G**), reflecting the failure of microglia to remove Homer1^+^ synapses. However, we found that the levels of active zone protein Bassoon (*34*, *35*) were significantly decreased in the 6 mo NL-F;Trem2 hippocampus (**Fig. 4A** and **B**). These data altogether suggest that that there may be exacerbated synaptopathology in the NL-F;TREM2 R47H mice.

**Fig. 4.**
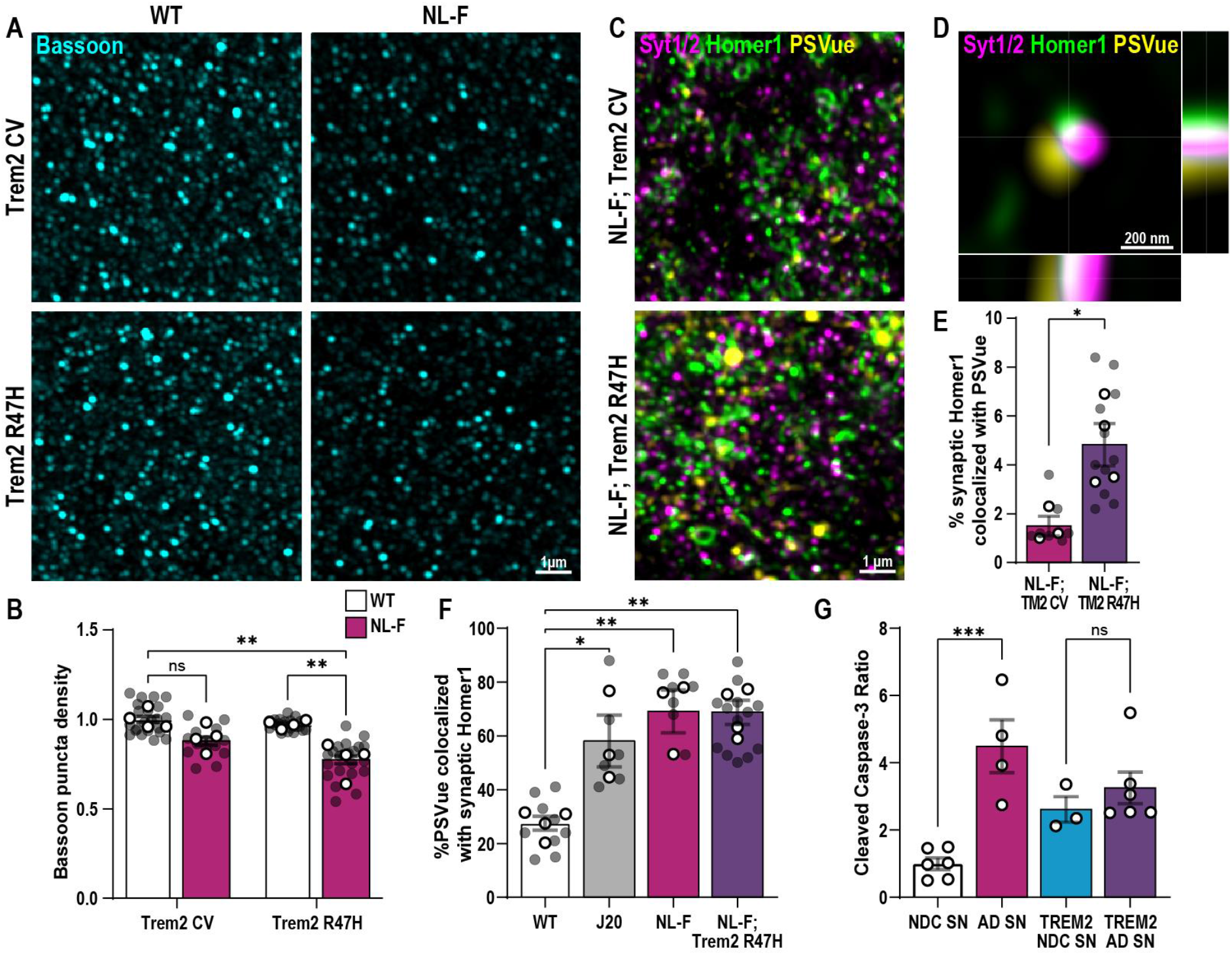
TREM2 loss-of-function leads to ePtdSer accumulation on AD synapses. **A)** SRM images from the hippocampal CA1 SR of 6 mo WT, Trem2 R47H KI, NL-F KI and NL-F KI; Trem2 R47H KI mice immunostained for Bassoon (magenta). **B)** Bassoon puncta density normalized to WT showing a decrease in NL-F KI; Trem2 R47H KI whereas that level of decrease was not yet seen at this early pre-plaque 6 mo NL-F hippocampus. **C)** Hippocampal dentate gyrus hilus of 6 mo NL-F KI and NL-F KI; Trem2 R47H KI mice ICV injected with PSVue 643 (yellow) and immunostained for Synaptotagmin1/2 and Homer1. **D)** Orthogonal view showing colocalized PSVue with synaptic markers. **E)** Percentage of synaptic Homer1 puncta within 0.25 μm of PSVue showing an increase in synaptic localized PSVue^+^ in NL-F KI; Trem2 R47H compared to NL-F KI. **F)** Percentage of PSVue within 0.25 μm of synaptic Homer1 puncta showing the specificity of synaptic PSVue in Aβ-driven AD models. **G)** Western blot densitometry showing the ratio of cleaved caspase-3/procaspase-3 in human NDC, AD, TREM2 NDC and TREM2 synaptosomes (SN) normalized to NDC SN. Data shown as mean ± SEM. Shaded points represent 1 ROI, open points represent animal average. 3 (B), 2-3 (E, F) ROIs per animal, n=3-4 animals, n=3-6 human cases (1 point represents 1 patient **(G)**). Two-way **(B)** or One-way **(F, G)** ANOVA followed by Bonferroni’s post-hoc test, unpaired t-test **(E)**, p-values shown as ns P>0.05; *P<0.05; **P<0.01; ***P<0.001. Scale bar 1 μm **(A, C),** and 200 nm **(D).**

To determine whether the failure by microglia to remove synapses leads to increased ePtdSer^+^ synapses in NL-F;Trem2 R47H KI brains, we performed intracerebroventricular (ICV) injection of PSVue to label ePtdSer on membranes of living mice (*10*). SRM imaging shows punctate PSVue in the hippocampus of NL-F and NL-F;Trem2 R47H mice, often co-localizing with markers immunoreactive for synaptic puncta (**Fig. 4C** and **D**). Importantly, a significantly higher percentage of synaptic Homer1 was labeled with PSVue in the NL-F;Trem2 R47H KI hippocampus as compared to those in the NL-F hippocampus (**Fig. 4E** and **fig. S8A**, **B**, **C** and **D**). These results, together with the *in vivo* microglia-synapse engulfment data (**Fig. 3C, D** and **E**), suggest that TREM2 is required for removal of synaptic ePtdSer in the pre-plaque NL-F brain.

In pre-plaque brains of mice carrying Aβ pathology, i.e., 6 mo NL-F, NL-F;Trem2 R47H KI and the 4 mo J20 transgenic mouse model of hAPP (*19*), we found higher percent of PSVue to co-localize with pre- and post-synaptic compartments versus healthy young adult WT mice (**Fig. 4F** and **fig. S8**). We also assessed for levels of apoptosis-like features in synaptosomes from patient brains. Direct PSVue labeling is not possible in frozen samples; as such, we probed for levels of caspase-3 activation, which is upstream of ePtdSer (*9*) and a marker for synaptosis (*12*). We found increased levels of caspase-3 activation in synaptosomes isolated from human AD brains as compared to those from NDC brains (**Fig. 4G** and **fig. S9**). Further, in a small cohort of patients carrying the AD-risk associated TREM2 variants, we found significantly higher levels of caspase-3 activation in their brain synaptosomes as compared to NDC, suggesting that TREM2 loss-of-function may exacerbate apoptotic features in human synapses (**Fig. S9**), as in mouse models. Altogether, these data in mice suggest that synaptic ePtdSer is a feature of AD synaptopathology that accumulates in AD brains, and that functional TREM2 is required to promote its removal by microglia.

Here, we show that synapses expose PtdSer upon Aβ challenge and that synaptic ePtdSer recruits microglia to phagocytose the synapse. Microglial removal of ePtdSer^+^ synapses aids in the resolution of oAβ-induced neuronal hyperactivity, suggesting that this form of synaptic regulation is not aberrant, at least at early stages of the pathology. Furthermore, we show that TREM2, a major risk factor for late-onset AD(*2*, *3*), mediates the selective engulfment of AD synapses by microglia. In accord, there are increased levels of apoptotic synapses remaining in the brains of AD humans and mouse models with dysfunctional TREM2. Our studies altogether suggest a role for ePtdSer-TREM2 in microglia-synapse engulfment in AD.

We propose a model in which oAβ induces ePtdSer on synapses that acts as ‘eat-me’ signal, rendering the ePtdSer^+^ synapses vulnerable to microglial engulfment. Whereas microglia can engulf ePtdSer^+^ neuronal cells in various injury and disease context (*36–38*), in the healthy normal brain, ePtdSer signals have been shown to occur on synaptic and other subcellular compartments of neurons to promote their phagocytosis in a context-dependent manner (*10, 11, 39*). In brains of AD mouse models and patients, local increase of caspase-3 activation and ePtdSer on hippocampal synapses occurs at the onset of synapse loss (our data here, *10*, *34–36*). ePtdSer could be induced on synapses in multiple ways: Endogenous Aβ production, as a natural by-product of activity-dependent APP cleavage, occurs on synaptic terminals (*43*). Inefficient clearing of Aβ from synapses can lead to its deposition and aggregation on synaptic membranes and/or disruption of synaptic dynamics including glutamatergic transmission and synaptic plasticity (*16*–*18*), altogether resulting in mitochondrial dysfunction and local activation of caspase-3, a well-established modulator of flippases and scramblases that regulate ePtdSer (*8*, *9*, *14*).

Here, we show that in pre-plaque AD brains, microglia, true to their role as brain-resident macrophages, sense synaptic ePtdSer for selective removal and resolution of oAβ-induced neuronal hyperactivity. The recognition of ePtdSer is mediated by TREM2 (*26*); in accord, we show that microglia with dysfunctional TREM2 fail to engulf AD ePtdSer^+^ synapses, resulting in an increased level of ePtdSer^+^ synapses remaining in the hippocampus along with significantly decreased levels of Bassoon, suggesting exacerbation of presynaptic defects in these pre-plaque brains. Whereas the role of TREM2 in plaque- and tangle-related neuropathology in AD has been widely studied (*44, 45*), little is known whether TREM2 impacts synapse engulfment in pre-plaque brains. TREM2 loss-of-function leads to decreased microglia-synapse engulfment across development and disease (our data here, *24*, *25*). Altogether, our data provide a possible explanation of how TREM2 variants may lead to increased risk for AD. Exact mechanisms of how TREM2 modulates synapse phagocytosis, for e.g., via through TREM2 shedding (*32*), and the functional consequences of the ePtdSer-TREM2 synapse engulfment dysregulation, particularly with aging, are yet to be determined.

Finally, we show that there is selectivity to which synapses microglia engulf in AD-relevant context. Our study provides a mechanistic insight into neuroimmune crosstalk at the synapse in AD-relevant context. Given that microglia are equipped with a multitude of phagocytic receptors and sensors that have been shown to directly or indirectly interact with ePtdSer (*8*), it is likely that TREM2 is not the sole mechanism by which ePtdSer is recognized and removed in AD brains (*36, 38*). An ensemble of ‘eat-me’ (*19*) and ‘don’t-eat-me’ (*46*) molecules on synapses may together determine the precision of which synapses are removed or not removed. Given the importance of synaptic homeostasis, especially as cells become senescent in the aging brain, what decisive and rapid alliance of various signaling molecules regulate microglia-synapse engulfment will be important to determine.

## Acknowledgments

We thank Dr. Frances Edwards (UCL) for provision of mice and PPL assistance; Dr. Samuel Barnes (UK DRI, ICL) for critical reading of this manuscript and advice on GCamp studies; members of the Hong lab and UK DRI at UCL for critical input to the manuscript, especially Gerard Crowley for surgical assistance; Dr. Ian White (UCL LMCB) for synaptosome EM preparation and imaging (Fig. S4A); Elena Ghirardello and Phillip Muckett for animal husbandry; Dr. Nicholas Cade for microscopy (UK DRI at UCL); and Dr. Andrey Klymchenko (Université de Strasbourg) for invaluable insight on ePtdSer visualization. The Queen Square Brain Bank is supported by the Reta Lila Weston Institute of Neurological Studies, UCL Queen Square Institute of Neurology.

## Licences

Ethics and guidelines for human tissue (18/LO/0721; IRAS Project ID: 246790) under UCL MTA 42-20. All animal experiments have been reviewed by UCL’s animal care committees and conducted in accordance with the regulations set out in the Animals in Scientific Procedures Act (ASPA) 1986. Animal experimental procedures were approved by the Institutional Animal Care and Use Committee of Seoul National University (approval ID no. SNU-191218-4-3, SNU-200904-2-4). All animal work was completed under Hong Lab PPL (PP1880415) and Edwards Lab PPL (70/8999) by licenced PIL holders.

## Authors’ contributions

Conceptualization: JRC, DS, SEL, SH. Data curation: JRC, DS, SEL, TC, NJ, SDS, JZG. Formal Analysis: JRC, DS, SEL, TC, NJ. Funding acquisition: SH, SEL, SC, OJF, DS. Investigation; JRC, DS, SEL, TC, NJ. Methodology; JRC, DS, SEL, SH, TC. Project administration; SH, BS, JH. Resources: JIL, CET, TL. Supervision: SH, SC, OJF, JRC. Validation: JRC, DS, SEL, TC. Visualization: JRC, DS, SEL, TC, NJ, SH. Writing – original draft: JRC, DS, SEL, SH.

## References and Notes

1. DeKosky, S.T. & Scheff, S.W. Annals of Neurology 27, 457–464 (1990).

2. Guerreiro, R. et al. New England Journal of Medicine 368, 117–127 (2013).

3. Jonsson, T. et al. New England Journal of Medicine 368, 107–116 (2013).

4. Podleśny-Drabiniok, A., Marcora, E. & Goate, A.M. Trends in Neurosciences 43, 965–979 (2020).

5. De Schepper, S., Crowley, G. & Hong, S. Developmental Neurobiology 81, 507–523 (2021).

6. Badimon, A. et al. Nature 586, 417–423 (2020).

7. Bartels, T., De Schepper, S. & Hong, S. Science 370, 66–69 (2020).

8. Lemke, G. Nature Reviews Immunology 19, 539–549 (2019).

9. Segawa, K. et al. Science 344, 1164–1168 (2014).

10. Scott-Hewitt, N. et al. The EMBO journal (2020).

11. Li, T. et al. The EMBO Journal 39, 1–20 (2020).

12. Ertürk, A., Wang, Y. & Sheng, M. Journal of Neuroscience 34, 1672–1688 (2014).

13. D’Amelio, M. et al. Nature Neuroscience 14, 69–79 (2011).

14. Sokolova, D., Childs, T. & Hong, S. Faculty Reviews 10, (2021).

15. Ebihara, T., Kawabata, I., Usui, S., Sobue, K. & Okabe, S. Journal of Neuroscience 23, 2170–2181 (2003).

16. Li, S. et al. Neuron 62, 788–801 (2009).

17. Renner, M. et al. Neuron 66, 739–754 (2010).

18. Zott, B. et al. Science 365, 559–565 (2019).

19. Hong, S. et al. Science 352, 712–716 (2016).

20. Hanshaw, R.G., Lakshmi, C., Lambert, T.N., Johnson, J.R. & Smith, B.D. ChemBioChem 6, 2214–2220 (2005).

21. Butovsky, O. et al. Nature Neuroscience 17, 131–143 (2014).

22. Gosselin, D. et al. Science 356, 1248–1259 (2017).

23. Sodero, A.O., Weissmann, C., Ledesma, M.D. & Dotti, C.G. Neurobiology of Aging 32, 1043–1053 (2011).

24. Krahling, S., Callahan, M.K., Williamson, P. & Schlegel, R.A. Cell Death and Differentiation 6, 183–189 (1999).

25. Lee, S.E. et al. Proceedings of the National Academy of Sciences of the United States of America 113, 6749–6754 (2016).

26. Wang, Y. et al. Cell 160, 1061–1071 (2015).

27. Keren-Shaul, H. et al. Cell 169, 1276–1290.e17 (2017).

28. Jaitin, D.A. et al. Cell 178, 686–698.e14 (2019).

29. Filipello, F. et al. Immunity 48, 979–991.e8 (2018).

30. Gratuze, M. et al. Journal of Clinical Investigation (2020).

31. Xiang, X. et al. Molecular Neurodegeneration 13, 1–14 (2018).

32. Kleinberger, G. et al. Science Translational Medicine 6, (2014).

33. Saito, T. et al. Nature Neuroscience 17, 661–663 (2014).

34. Altrock, W.D. et al. Neuron 37, 787–800 (2003).

35. Lazarevic, V., Schöne, C., Heine, M., Gundelfinger, E.D. & Fejtova, A. Journal of Neuroscience 31, 10189–10200 (2011).

36. Brelstaff, J., Tolkovsky, A.M., Ghetti, B., Goedert, M. & Spillantini, M.G. Cell Reports 24, 1939–1948.e4 (2018).

37. Garcia-Reitboeck, P. et al. Cell Reports 24, 2300–2311 (2018).

38. Tufail, Y. et al. Neuron 93, 574–586.e8 (2017).

39. Ruggiero, L., Connor, M.P., Chen, J., Langen, R. & Finnemann, S.C. Proceedings of the National Academy of Sciences of the United States of America 109, 8145–8148 (2012).

40. Phongpreecha, T. et al. Science Advances 7, 1–15 (2021).

41. Mattson, M.P., Keller, J.N. & Begley, J.G. Experimental Neurology 153, 35–48 (1998).

42. Park, G. et al. Cell Reports 31, 107839 (2020).

43. Cirrito, J.R. et al. Neuron 58, 42–51 (2008).

44. Lee, S.H. et al. Neuron 109, 1283–1301.e6 (2021).

45. Deczkowska, A., Weiner, A. & Amit, I. Cell 181, 1207–1217 (2020).

46. Lehrman, E.K. et al. Neuron 100, 120–134.e6 (2018).

