## Supplementary Materials for "Microglia Detect Externalized Phosphatidylserine on Synapses for Elimination via TREM2 in Alzheimer’s Disease Models"

Materials and Methods
Figs. S1 to S9
Tables S1 to S3
Captions for Movies S1 to S5
Movies S1 to S5

### Materials and Methods

#### Animals

All experiments have been reviewed by UCL's animal care committees and conducted in accordance with the regulations set out in the Animals in Scientific Procedures Act (ASPA) 1986. Sprague Dawley rats obtained from Charles River UK and Homer1-eGFP (*15*) obtained from Japan (kind gift from S. Okabe) were used for primary neuronal cultures at embryonic day (E18). C57BL6/J (WT) mice obtained from Charles River UK and Trem2 R47H KI mice (imported from JAX, C57BL/6J-*Trem2*$^{em1Adiuj/J}$ Strain #027918) were used for primary microglial culture preparation at P0-P4. *App*$^{NL-F}$ KI mice (*33*) (kindly provided by Takaomi Saido, Riken and distributed by Frances Edwards, UCL) were crossed to Trem2 R47H KI mice for *in vivo* labelling of lipids, synapse loss and microglial engulfment studies. APP transgenic J20 mice (kindly provided by Lennart Mucke and distributed by Patricia Salinas, UCL) were used at age 3 months for *in vivo* labelling of lipids. For all experiments appropriate sex- and age-matched controls were used. For *App*$^{NL-F}$ KI and Trem2 R47H KI genotype, homozygous mice were used, for Homer1-eGFP heterozygous mice were used.

#### Primary Neuronal Culture

Primary hippocampal neurons were prepared as previously described (*25*) from E18 Sprague Dawley rats and Homer1-eGFP mice of either sex (n=10-15 pups per preparation). Briefly, hippocampi were dissected, dissociated with papain, and triturated with a polished half-bore pasteur pipette. Next, cells were resuspended in Hank's Balanced Salt Solution (HBSS; HyClone, Logan, UT) supplemented with 0.6 % glucose, 1 mM pyruvate, 2 mM GlutaMAX (Gibco), and 10 % (v/v) FBS (HyClone) and plated on Poly-D-lysine (PDL)-coated glass coverslips in a 60-mm Petri dish or 35 mm glass-bottom culture dish (81158, ibidi). 4 h after plating, the medium was replaced with neurobasal medium (Invitrogen) supplemented with 2 % (v/v) B-27 (Invitrogen), 0.5 mM GlutaMAX. Half of the medium was replaced by a new neurobasal media with B-27 and L-glutamine at DIV 4, 7 and 14. 4 mM 1-β-D-cytosine-arabinofuranoside (Ara-C; Sigma) was added as needed. Cells were maintained in an incubator at 37°C and 5% $CO_2$ and used at DIV 17-21 for experimental procedures.

#### Primary Neuron jGCaMP7 Transfection

Neurons were transfected using a modified calcium-phosphate method as previously described (*25*) using a pAAV-syn-jGCaMP7c-WPRE plasmid (Addgene). Briefly, 6 µg of DNA and 9.3 µl of 2 M $CaCl_2$ were mixed in distilled water to a total volume of 75 µl and the same volume of 2x BBS [50 mM BES, 280 mM NaCl, and 1.5 mM $Na_2HPO_4$ (pH 7.1)] was added. The cell culture medium was completely replaced by transfection medium (MEM; 1 mM sodium pyruvate, 0.6% glucose, 10 mM HEPES, 1 mM Kynurenic acid, and 10 mM $MgCl_2$, pH 7.71), and the DNA mixture was added to the cells and incubated in a 5% $CO_2$ incubator for 60 min. Cells were washed with a washing medium (pH 7.30), and then returned to the original culture medium. Neurons were transfected at DIV 8-9 and analyzed at DIV 16-21. pAAV-syn-jGCaMP7c-WPRE plasmid was purchased from Addgene.

#### Primary microglial culture preparation

Primary mouse WT and Trem2 R47H KI microglial cultures were prepared at P0-P4 mice from either sex (n=8-10 pups per preparation). Mouse brains were dissected in cold HBSS on ice and the cortices and hippocampi were isolated. Tissue was homogenized with 2 ml stripette (15 strokes). Next, the homogenate was put through a pre- wet 70 μM strainer and centrifuged at 400g for 5 min at 4°C. The supernatant was removed and the cell pellet was resuspended in ice cold 35% isotonic percoll. The interface was carefully created with HBSS. The samples were centrifuged for 40 min at 4°C at 2800g with no break and with slow acceleration and deceleration. The myelin layer and supernatant was aspirated and the cell pellet was washed in HBSS. The cells were centrifuged for 5 min at 4°C and 400 g. The supernatant was removed and cells were resuspended in 1ml microglial media (DMEM F12 Gibco, 5% fetal bovine serum Gibco, 1% pen-strep Gibco, 50 ng/mL CSF1 416-ML-010/CF RnD Systems, 50 ng/mL TGFb1 7666-MB-005/CF RnD Systems, 100 ng/mL CX3CL1 472-FF-025/CF RnD Systems) for cell counting. Cells were plated in borate buffer 0.1 M pH 8.5 PDL (Gibco) coated 12-well plates (CC7682-7512, Starlab) in 1 ml of microglial media at a density of 650,000 per well. Cells were maintained in an incubator at 37°C and 5% $CO_2$. 90% of media was changed the day after and subsequently half of the media was changed every 2 days.

Primary microglial cells are supplemented with TGFβ, which has been shown to imprint key microglial signature genes such as *Tmem119* (*21*, *22*). Additionally, mCSF is added to the media to stimulate microglial survival and fractalkine as the ligand for the key microglial homeostatic receptor, CX3CR1. The combination of TGFβ, mCSF and fractalkine is used to mimic a more homeostatic *in vivo*-like profile of microglia as shown by high levels of *Tmem119* mRNA in *Cx3cr1*$^{+}$*Trem2*$^{+}$ *Itgam*$^{+}$ primary microglial cells (**Fig. S3**).

**RNA isolation, reverse transcription and RT-qPCR**

Primary microglial cells were lysed and scraped off using TRIzol reagent (15596026, Invitrogen) after which chloroform was added to separate the homogenate layers. RNA was precipitated from the aqueous layer using 2-propanolol and then washed with ethanol. The RNA pellet was re-suspended in nucleus-free-water after which RNA purity and concentration was assessed by Nanodrop. mRNA was converted to cDNA using the qScript cDNA SuperMix reverse transcription kit as described by the manufacturer (95048, Quantabio). For RT-qPCR, 12ng of cDNA was loaded in triplicates per gene in a total volume of 20 μl using the SYBR green PCR master mix as described by the manufacturer (4309155, ThermoFisher). The reaction was run using a LightCycler 96 Instrument (Roche) with white 96-well plates (04729692001, Roche). Triplicate Ct values were averaged and data is shown as respective to the geomean of 3 housekeeping genes (*Actb*, *Gapdh*, *Rpl32*) using the Ct delta method ($2^{-\Delta\Delta Ct}$). Primers purchased from IDT were used at a concentration of 200nM, see **Table S1** for sequences.

**Neuron-microglia co-culture**

Primary microglia at DIV 7 were detached with ice-cold PBS and centrifuged for 5 min at 4°C and 400 g. The supernatant was removed and cells were resuspended in 1 ml neuron culture media supplemented with CSF1, TGFβ1 and CX3CL1. Cells were plated on the DIV 14 neurons at the ratio of 2:1. Cells were co-incubated in an incubator at 37°C and 5% $CO_2$ for 7 days before analysis.

**Neuronal Calcium Imaging**

To measure neuronal spontaneous calcium transients, jGCaMP7 expressing primary neurons cultured with or without microglia were assessed using a spinning disk confocal microscope (ECLIPSE Ti-E, Nikon) with a Plan Apo 60×/NA 1.40 oil objective, and a Neo sCMOS camera

(Andor Technology) at 37°C. Time-lapse images were acquired every 100 ms for 1 min. First, regions of interest (ROIs) were manually drawn around individual dendritic spines, and then relative fluorescence change (ΔF/F) versus time was measured for each ROI. Spontaneous calcium transients were identified as changes in ΔF/F that were larger than 10% of the baseline intensity and the high-frequency was defined if showed the calcium transient frequency > mean + 2 SD of the initial value.

**Live labelling**

For live cell labelling of externalized phosphatidylserine (ePtdSer), 1 mM PSVue® 550 (P-1005, Molecular Targeting Technologies; prepared following manufacturer's instructions) was diluted in Tyrode's solution (136 mM NaCl, 2.5 mM KCl, 2 mM $CaCl_2$, 1.3 mM $MgCl_2$, 10 mM HEPES and 10 mM Glucose, pH 7.4) at 1:1000 and incubated for 10 min before imaging. For live-cell microglial labelling, fluorescent conjugated plant lectin Griffonia (Bandeiraea) simplicifolia lectin I, Isolectin GS-$IB_4$-647 (IB4) (I32450, ThermoFisher) was diluted in Tyrode's solution at 1/1000 and incubated for 10 min before imaging. IB4 binds selectively to microglial RET receptor tyrosine kinase and is commonly used as a microglial marker in the brain.

**High-resolution cell imaging and analysis**

Airyscan live-cell images were acquired with a laser scanning LSM880 Airyscan microscope, using a Plan-APO 20X/NA 0.8 objective (Zeiss). Emission filter bandwidths and sequential scanning acquisition were set up, to avoid any possible spectral overlap between fluorophores with 37°C and 5% $CO_2$ maintained. 3D time-lapse images were acquired in z-stack step size 800 nm X 15 steps every 2 min for 1 h and subsequently processed using Imaris software (Bitplane). For Homer1-eGFP and PSVue co-localization analysis, z-stack images were acquired on a LSM880 Airyscan microscope using 63×/NA 1.40 objective with 0.3 µm z-steps. The percentage of Homer1 and PSVue co-localization was calculated for every z-step using Fiji (NIH software). For quantification of preferential contact and engulfment of Homer1-eGFP$^+$ PSVue$^+$ dendritic spines by microglia, ROIs with mobile Homer1-eGFP puncta within 5 µm from microglial were selected and the ratio of colocalization with PSVue was analyzed.

For quantification of PSVue level post fixation of neuron-only and co-cultured neurons with microglia in GCaMP7 experiments, z-stack images were acquired on a LSM980 Airyscan microscope using 63×/NA 1.40 objective with 0.17 µm z-steps. The puncta number per 100 $\mu m^2$ and mean intensity of PSVue were measured per ROI using Fiji (NIH software).

**Intracerebroventricular PSVue injection**

For *in vivo* labelling of ePtdSer, 1 mM PSVue® 643 (P-1006, Molecular Targeting Technologies) was used following manufacturer's recommendations. 4-months-old WT and J20 Tg animals, 6-month-old *App*$^{NL\text{-}F}$ KI and *App*$^{NL\text{-}F}$ KI; Trem2 R47H KI mice littermates were used. Mice were anaesthetized with 4% inhaled Isoflurane (Forane, Abbott Laboratories) and placed in a stereotaxic apparatus (504926, World Precision Instruments Ltd). Anaesthesia was maintained at 1.5% in 250 ml/min oxygen flow. Under aseptic conditions, a midline incision was made to reveal the skull. Two holes were drilled in the skull using a 0.8 mm diameter burr (503599, OmniDrill35 Micro Drill, World Precision Instruments Ltd) to allow for a bilateral injection into the lateral ventricles. Next, 1.5 µl of sterile PSVue was injected using a 10 µl syringe with a fine borosilicate glass capillary (Hamilton) in the following coordinates: 0.5 mm anterior/posterior, +/-1.0 mm lateral, and -2.3 mm dorsal/ventral from bregma (Paxinos and Franklin's The Mouse Brain in Stereotaxic Coordinates, Fourth Edition). Infusion was performed with a Microinjection Syringe Pump (World Precision Instruments Ltd) at a rate of 0.3 µl/min. The needle was kept in this position for an additional 5 min after injection and then retrieved slowly to avoid backflow. The incision on the scalp was closed with Vetbond tissue adhesive (3 M). Subcutaneous carprofen (Carprieve, 5 mg/g body weight) and buprenorphine (Vetergesic, 0.1 mg/g body weight) diluted in 0.9% saline were administered peri-operatively. 24 h after injection, the animals were perfused with 4% PFA for histological analysis.

**Crude synaptosome preparation**

Synaptosomes were prepared from fresh mouse and frozen human postmortem brain tissue provided by the QSBB, (see **Table S2** for patient information). WT mice aged 2-4 months were used (3-5 animals per preparation). In brief, mice were intra-cardiac perfused with 10ml cold PBS. The hippocampi and cortices were dissected on ice. For post-mortem human tissue, synaptosomes were prepared from the frontal cortex. Synaptosomes were biochemically isolated as previously described (*23*). Tissue was weighed and homogenized in 5 volumes of sucrose homogenization buffer (5 mM HEPES pH 7.4, 320 mM sucrose, 1 mM EDTA) using a Dounce homogenizer with 15-20 strokes. The homogenate was centrifuged at 3,000 g for 10 min at 4°C and the supernatant was saved as total homogenate fraction (THF). The THF was centrifuged again at 14,000 g for 12 min at 4°C and supernatant was saved as cytosolic fraction. The pellet was carefully resuspended in 550 µl of Krebs-Ringer buffer (KRB: 10 mM HEPES, pH 7.4, 140 mM NaCl, 5 mM KCl, 5 mM glucose, 1 mM EDTA) and 450 µl of Percoll solution (for a final concentration of 45%). The solution was mixed by gently inverting the tube and an interface was slowly created with KRB. After centrifugation at 14,000 g for 2 min at 4°C, the synaptosomal fraction was recovered at the surface of the flotation gradient and carefully re-suspended in 1 ml of KRB to wash. The synaptosomal preparation was centrifuged at 14,000 g for 1 min at 4°C, after which the pellet was re-suspended in KRB. When done on fresh tissue, this protocol yields synaptosomes that are electrically functional for several hours post isolation as they can be depolarized and stimulated with KCl and NMDA respectively. After isolation, a standard BCA protein assay was performed to obtain amount of protein for subsequent assays.

**Synthetic humanised oAβ 40-S26C dimer treatment**

Primary cultures or fresh synaptosomes were treated with 50 nM oAβ 40-S26C dimer (018-71, Phoenix) vs PBS control for 1 h in an incubator at 37°C and 5% $CO_2$. Experimental procedures were either performed on live cells during the 1h window or on fixed cells post treatment. Fresh mouse synaptosomes were immediately divided into Eppendorfs at 2-2.5 mg of protein and re-suspended in total 1 ml KRB in 50 nM of oAβ 40-S26C dimer or just buffer and PBS as control and left overnight at 4°C on nutator. Synaptosomes were then centrifuged at 14,000 g for 1 min at 4°C, supernatant was discarded and synaptosomes were washed in 1ml PBS, after which they were centrifuged at 14,000 g for 1 min at 4°C to obtain oAβ-synaptosomes and control-synaptosomes.

**Immunocytochemistry (ICC)**

Cells and synaptosomes were fixed for 10 min at room temperature (RT) in 4% (w/v) PFA, 4% (w/v) sucrose in PBS, pH 7.4 and subsequently permeabilized with 0.25% Triton X-100 in PBS for 3 min at RT. The cells were then blocked for 1 h at RT in 10% (w/v) Bovine serum albumin (BSA). Cells and synaptosomes were incubated at 4°C overnight in primary antibodies (1/1000) after which the cells were washed in PBS and incubated with secondary antibodies (1/1000) for 1h at RT (see **Table S3** for antibody information). The immunostaining of synaptosomes was performed in Eppendorfs with a centrifugation step at 14,000 g for 1 min at every wash step. At the end of the protocol, synaptosomes were re-suspended in pro-long gold antifade mounting media (P36930, Invitrogen) and put through a 1 ml insulin syringe to further homogenize synaptosomes. This solution was then mounted on glass slides (SuperFrost GOLD Adhesion Slides 11976299, Fisher Scientific).

For fresh synaptosome labelling of ePtdSer, synaptosomes were left on nutator at RT in PSVue 643 for 1 h after which excess PSVue was centrifuged at 14,000 g for 1 min and washed. Synaptosomes were then fixed for ICC.

**Synaptosomes conjugation to pHrodo amine-reactive labels**

Synaptosomes were conjugated to low background pH-sensitive dyes, which fluoresce brightly upon acidification (pH 4-6) such as in late endosomes and lysosomes using an adapted protocol from (*46*). pHrodo dyes are photostable allowing for multicolor and long-term imaging. In brief, pHrodo™ Red, succinimidyl ester (P36600) and pHrodo™ Deep Red Antibody Labeling Kit (P35355) were dissolved as described in the manual. Human post-mortem synaptosomes were conjugated to pHrodo red whereas mouse synaptosomes were conjugated to both pHrodo red and deep red for preferential engulfment studies. 1 mg of synaptosomes were left at RT on nutator for 2 h in sodium bicarbonate 0.1 M with respective pHrodo at a concentration of 1 mg/ml. After conjugation, synaptosomes were centrifuged at 14,000 g for 1 min, then washed with 1 ml PBS and centrifuged again at 14,000 g for 1 min. Synaptosomes were then resuspended as described by manufacturer. Next a standard BCA protein assay was performed on pHrodo conjugated synaptosomes to quantify protein concentrations for subsequent engulfment assays. Synaptosomes were then aliquoted respectively and stored at -80°C. Prior to engulfment analysis, the degree of pHrodo labelling (DOL) was assessed as described by manufacturer. In brief, synaptosomes were mixed 1:3 in PBS pH 2 to activate the fluorophores. The relative efficiency of the labeling reaction was determined by measuring the absorbance of the protein at 280 nm and the absorbance of the dye at its excitation maximum using a UV-Vis Spectrophotometer. This was done as a control to ensure that the DOL was similar between pHrodo and treatment paradigms.

**Synaptosome electron microscopy**

Synaptosome pellet was pooled from the hippocampi and cortices of 3 wild-type mice and prepared as described above. At all solution changes the pellet was resuspended in the new solution, and centrifuged to re-pellet the sample before removal of that solution and replacement and resuspension with the next. The pellet was fixed in 2% glutaraldehyde/1.5% formaldehyde in 0.1 M sodium cacodylate for 30 min at room temperature. The fix was replaced with 1% osmium tetroxide/1.5% potassium ferricyanide for 1 h at 4°C. Post osmium treatment the sample was washed three times in cacodylate buffer then incubated in 0.1% tannic acid in 0.05M cacodylate for 40 min at room temperature. The sample was washed twice in 0.5M cacodylate and once in dH2O before dehydration in 70%, 90% and 2 x 100% ethanol each for 10 min. Samples were embedded in resin beginning with 1:1 mix of propylene oxide:epon resin for 60 min, then 100% epon overnight and 4 h in fresh epon the next day before being transferred to a 60°C oven to polymerize overnight. Ultrathin sections of the sample at 70 nm thickness were cut using a diamond ultra 45° knife (Diatome) on a Leica UC7 ultramicrotome, and collected on formvar coated copper 2 × 1mm slot grids. Sections were further contrast stained with Reynolds Lead citrate before being imaged on a transmission electron microscope (T12 Tecnai Spirit biotwin, FEI) equipped with a charge-coupled device camera (SIS Morada; Olympus).

#### *In vitro* microglia-synaptosome engulfment assay

For preferential *in vitro* engulfment assays using mouse synaptosomes, primary mouse microglia were treated with oAβ-synaptosomes and control synaptosomes. 1 μg of both control and oAβ-synaptosomes were added to the same well in microglial media. For engulfment assays using human post-mortem synaptosomes, primary mouse microglia were either treated with 1μg of Alzheimer's disease (AD) or non-demented control (NDC) synaptosomes in separate wells. Plates were then placed in a cell discoverer 7 (CD7) with the incubator at 37°C and 5% $CO_2$. Fluorescent (594 nm and 647 nm) and brightfield (oblique and phase) images were acquired at a x20 objective (x0.5) at intervals of 2-5 min. Laser settings were set to the same settings for both red and deep red pHrodo. A 3-slice z-stack was taken at 1.5μm interval to ensure that imaging was within focus throughout the imaging session however, one plane was used for analysis. 2 ROIs per well were taken with an average of 40 cells per ROI. An imaging session lasted 0-15 h, whereby plateau was reached within the first 6 hr, with t=0 being the addition of synaptosomes. The plateau phase persisted and a decrease in the pHrodo signal was only observed after 48-72 h. Background subtraction was performed on ImageJ for red and deep red pHrodo at 1 pixel. For analysis, a plug-in on ImageJ was used, z-profile axis, which measures intensity of a given channel with respect to time. Fluorescence intensity at t=0 was subtracted from subsequent time frames. Experimental replicates were analyzed separately. Data was either shown as fluorescence intensity with time, engulfment (area under curve (AUC) at 50% of the peak pHrodo fluorescence intensity) or engulfment ratio. AUC at 50% peak pHrodo intensity for all experiments was approximately at 3 hr. For AUC at 50%, pHrodo intensity was normalized to average control. For engulfment ratio, the sum of control and oAβ-synaptosome fluorescence intensity was added per well and then individually divided by this total sum, to obtain a fraction per well.

#### Annexin-V treatment

Synaptosomes and neuron-microglia co-cultures were treated with Annexin-V (AnnxV), a protein that specifically binds to ePtdSer with high affinity, which has been used to mask PtdSer on apoptotic cells to block macrophage phagocytosis *(24)*. Synaptosomes were pre-treated with either buffer, 1 µg/mL or 10 µg/mL of purified recombinant AnnxV (556416, BD BioSciences, 0.5mg/mL stock concentration) in 100 µl 1x AnnxV binding buffer (556454, BD Biosciences), 0.1 M HEPES (pH 7.4) 1.4 M NaCl, 25 mM $CaCl_2$ (AnnxV binding to ePtdSer is calcium-dependent) for 1 h at RT. These concentrations have been previously used in literature and suggested by the manufacturer (5-15μg). We ensured that microglia were not exposed to AnnxV by washing the synaptosomes to remove excess AnnxV prior to application. Synaptosomes were centrifuged at 14,000 g for 1 min to remove excess AnnxV and re-suspended in 1x AnnxV binding buffer. Control and oAβ-synaptosomes were pre-treated simultaneously in the same Eppendorf, whereas NDC and AD human synaptosomes were pre-treated separately. Neuron-microglia co-cultures were treated with 0.1 ug/mL of AnnxV simultaneously with Aβ treatment.

#### Bafilomycin A1 treatment

Post microglia-synaptosome engulfment assays, 50 nM bafilomycin A1 (B1793, Sigma-Aldrich), which increases lysosomal pH to 6, was added to microglial wells. Imaging and analysis were performed as described above. Background subtraction was performed at all time-points using the intensity at t=0 (when synaptosomes were added). Data shown as t=0 when bafilomycin treatment was added.

**Western-blotting**
A BCA protein assay was used to determine the amount of protein in fractions used for western-blot analysis. 20-40 µg of protein were loaded either on 4-12% Bis-Tris gels or 10% Tris-Glycine gels (Thermofisher Scientific). Gels were run with appropriate sample buffer and running buffer. Gels were transferred to nitrocellulose membranes using an iBlot 2 Dry Blotting System (Thermofisher Scientific) as described by the manufacturer. Prior to blocking, Aβ blots were left in PBS and microwaved at 80% strength for 1 min 30s on each side, allowing for 3 min 30 s to settle after each side to expose the epitope. Blots were blocked in casein PBS (1:1, Biorad) for 30 min at RT on shaker and then probed overnight at 4°C on shaker with primary antibodies (1:1000) in casein PBS-T (0.01%). Blots were washed then probed with secondary antibodies in casein PBS-T (0.01%) for 1 h at RT on shaker then visualized either fluorescently on a ChemiDoC (BioRad) system or with HRP-substrate on an Amersham Imager 680 (Bioke) system.

**Immunohistochemistry (IHC)**
IHC was performed on brain slices from mice intracardiac perfused with 4% PFA and post-fixed for 24 h. 30 µm free-floating tissue sections were washed in PBS on a perturbator followed by pre-treatment in 1% Triton X-100 in PBS for 20 min, rinsed in PBS for 5 min, and treated with 0.3% Triton X-100. Blocking solution and primary and secondary antibody mixtures were centrifuged at 17,000 g for 5 min just before use. Sections were then blocked in 20% NGS, 1 % BSA, and 0.3 % Triton in PBS for synapse IHC or in 10% NGS, 10% FBS, 1 % BSA, and 0.3 % Triton in PBS for microglial engulfment IHC for 2 h followed by primary antibody incubation overnight at 4℃. Sections were washed in PBS for 10 min, 0.3% Triton X-100 in PBS for 30 min, followed by secondary antibody incubation for 4 h at room temperature. See **Table S3** for antibody information. Sections were then washed in PBS for 10 min, incubated in 1:10000 DAPI in PBS for 10 min and washed in 0.3 % Triton for 15 min. Finally, sections were mounted onto slides with ProLong Glass mounting medium and procured for at least 48 h before imaging. IHC of PSVue-labelled tissue was as described above, with the following substitutions: TBS instead of PBS; Alexa Fluor 546 goat anti-rabbit instead of Alexa Fluor 594 goat anti-rabbit.

**Super-resolution imaging and analysis**
Super-resolution synapse images were acquired on a Zeiss LSM 880 microscope with Airyscan detector using a 63x, 1.4NA oil immersion Plan-Apochromat objective (theoretical maximum resolution: 140 nm lateral, 350 nm axial). A zoom factor of 1.8x and frame size of 2048x2048 was used for all images, resulting in an XY pixel size of 37 nm. Z-step size was 144 nm, with 8 steps per Z-stack, resulting in a stack thickness of 1.15 µm. Scan speed was 5, line averaging 2, gain 800 and digital gain 1. A laser power of ~0.15% and ~0.45% was used for the 488 nm and 594 nm lasers, respectively. The Airyscan detector was aligned before imaging each new slide. Three regions of interest were acquired in the center of the hippocampal CA1 stratum radiatum for each brain section. Super-resolution synapse images were processed in Zen Black using 3D Airyscan Processing at strength 6.0. Images exhibiting drift or anomalous staining were excluded from analysis. Imaris software was used for pre- and postsynaptic puncta detection. Channel brightness was adjusted to maximize the visualization of immunoreactive puncta. The spots detection

function was used to generate spots, with Region Growing, Shortest Distance Calculation and Background Subtraction enabled. Spots size was selected to maximize the detection of immunoreactive puncta (XY diameter 0.15 µm, Z diameter 0.45 µm, for both Homer1 and Synaptotagmin1/2 puncta). Spots were classified using the 'Intensity Center' filter. The threshold value was selected to maximize the detection of immunoreactive puncta. The threshold value varied between channels and immunostainings but was kept consistent between pairs of images that were compared. Local Contrast was used to define the spot growth boundary, using a value that appropriately reflected the immunoreactive signal of the source channel. A volume filter was then applied to remove spots smaller than 0.005 $\mu m^3$. Pre- and postsynaptic spots were colocalized using a MATLAB colocalization script (Colocalize Spots XTension), using a colocalization distance of 0.25 µm between spot centers. Synapse density is shown as $App^{NL\text{-}F}$ KI mice normalized to WT mice, and $App^{NL\text{-}F}$ KI x *Trem2* R47H KI normalized to *Trem2* R47H KI mice.

PSVue-labeled synapse images were acquired on a Leica Stellaris STED microscope using a 100x, 1.4NA oil immersion Plan-Apochromat objective. A zoom factor of 2x and frame size of 2048x2048 was used for all images, resulting in an XY pixel size of 29 nm. Z-step size was 200 nm, with 11 steps per Z-stack, resulting in a stack thickness of 2.2 um. Images were acquired in Photon Counting mode with a scan speed of 600 Hz and line accumulation 8. A laser power of 5%, 5% and 10% was used for the 499 nm, 557 nm and 653 nm lasers, respectively. Three regions of interest were acquired in the center of the hippocampal dentate gyrus hilus for each brain section. Confocal images of PSVue-labelled synapses were deconvolved using Leica's LIGHTNING deconvolution. Imaris was used for pre- and postsynaptic puncta detection as described above. The Surfaces function was then used to generate volumes representing the PSVue signal. Pre- and postsynaptic spots within 0.25 µm of a PSVue surface were determined using a MATLAB colocalization script (Spots Close To Surface XTension).

***In vivo* microglial engulfment imaging and analysis**

Images were acquired on a Zeiss LSM 800 confocal microscope using 63x magnification. Frame size of 2048 x 2048 was used for all images. 11 µm Z-stack was acquired with a voxel size of 0.05 x 0.05 x 0.220 $\mu m^3$. Three regions of interest were acquired in the hippocampal CA1 stratum radiatum per section and three sections were analyzed per brain. Images were first processed with ImageJ using background subtraction. Imaris 3D surface rendering was used to first create a surface on the P2Y12 channel. Subsequently, CD68 channel was masked to the P2Y12 surface to obtain only CD68 within the analyzed cell. A surface was created on the CD68 masked channel and a volume filter of > 0.01 $\mu m^3$ was applied. Homer1 channel was then masked to the CD68 surface to obtain Homer1 inside $CD68^+$ vesicles, and a surface was created on thus masked Homer1 channel. Homer1 surfaces of volume smaller than 0.001 $\mu m^3$ were filtered out and not included in the analysis. Volume of thus obtained Homer1-immunoreactive puncta was then normalized per cell volume to account for variance due to difference in cell size. Imaris 3D surface function algorithm settings were the following for all channels: "Shortest distance calculation", "Absolute intensity", and grain size of 0.1 µm was used. The threshold values were adjusted to optimize visualization of individual channels, and varied between channels, but were kept constant within a channel for all images analyzed. Data shown as engulfment index: (volume of Homer1 in CD68 /microglial cell volume) *100.

**Statistics**

Statistical tests were performed using GraphPad Prism 9.0 (GraphPad Software) as appropriate. For multiple comparisons, Tukey or Bonferroni's post hoc test were used. Data points shown as either ROIs or average per animal. Statistical tests have been performed per animal for animal studies and per ROI for cell culture experiments. Data shown as mean ± SEM or SD. P-values shown as ns $P>0.05$; *$P<0.05$; **$P<0.01$; ***$P<0.001$; ****$P<0.0001$.

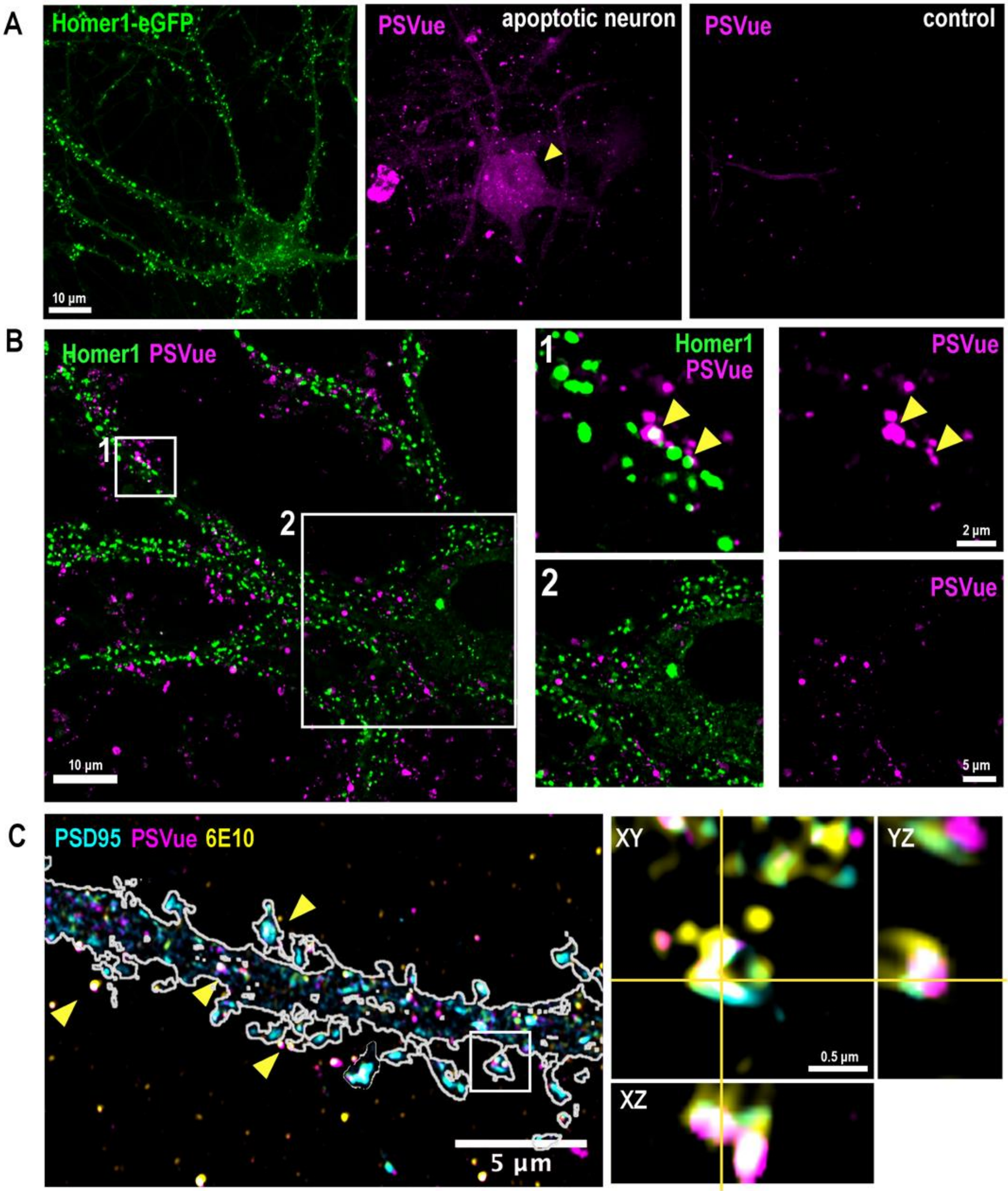


**Fig. S1. oAβ induces focal PtdSer externalization on dendritic spines.**
**A**) Representative image of healthy Homer1-eGFP neuron (green, left panel), apoptotic neuron as positive control for PSVue 550 (magenta, middle panel) staining across whole neuron and PSVue staining on healthy neuron (right panel). **B)** SRM images of neuronal PSvue staining 1 h after 50 nM oAβ treatment. Note PSVue staining in inset 1 and 2 is localised to Homer1-eGFP dendritic spines as indicated by yellow arrowheads and does not fluoresce across whole neuron. Refer to **(A)** for positive control of apoptotic neuron. **C**) SRM images of PSD95 (cyan) and PSVue and 6E10 (yellow) 1 h after oAβ treatment. Yellow arrowheads indicate colocalized PSD95, PSVue and 6E10 on dendrites. Inset shows orthogonal view of a single spine with PSD95, PSVue and 6E10 colocalisation. Scale bar 10 µm **(A)**, 10, 2 and 5 µm **(B)** 5 and 0.5 µm **(C)**.

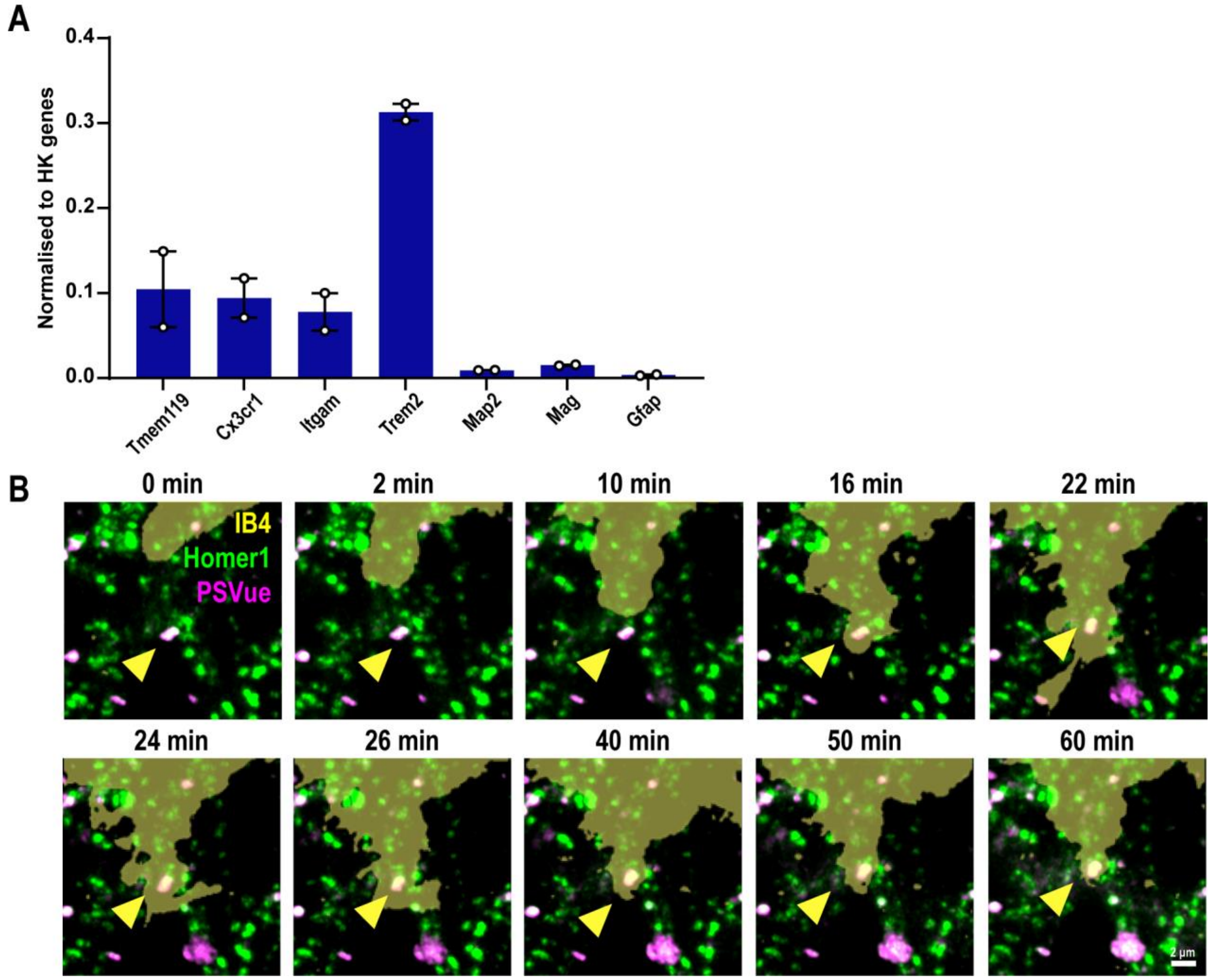


**Fig. S2. Primary microglia contact ePtdSer$^+$ dendritic spines.**

**A)** Primary microglia grown with TGFβ express high mRNA levels of microglial genes including *Cx3cr1*, *Itgam*, *Trem2* but also homeostatic *Tmem119* with little contamination from neurons (*Map2*), astrocytes (*Gfap*), and oligodendrocytes (*Mag*). Gene expression shown as normalized to the geomean of 3 house-keeping genes (*Actb*, *Gapdh* and *Rpl32*). **B)** Time-lapse sequence images of 10 consecutive optical sections ($\Delta z = 0.4$ μm) showing primary Homer1-eGFP neurons (green), PSVue (magenta) and microglia (yellow, labelled with Isolectin B4-647, IB4-647). Yellow arrowheads indicate PSVue$^+$ Homer1-eGFP dendritic spines contacted by microglia over 1 h. See Supplemental movie 1. 1 point represents 1 well, n=2 wells. Scale bar 2 μm.

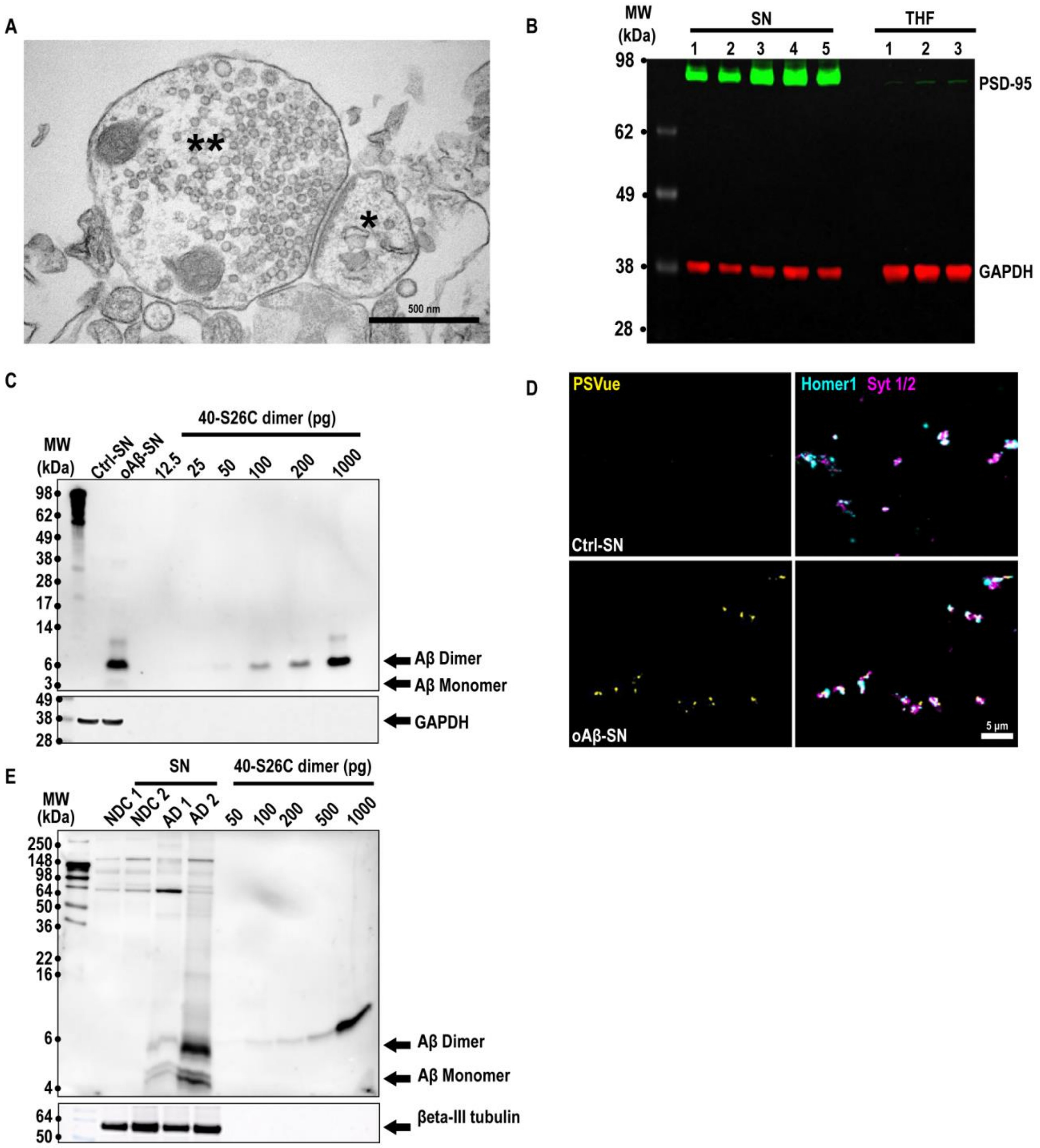


**Fig. S3. oAβ induces ePtdSer on fresh mouse synaptosomes.**
**A)** Crude synaptosomes were prepared using a protocol that yields electrically functional synaptosomes for several house post isolation, which can be depolarized and stimulated with KCl and NMDA respectively. Electron microscopy of crude synaptosome preparation showing intact pre- (two asterisks) and post-synaptic sites (one asterisks). **B)** Western blot showing enrichment of synaptic markers PSD-95 (green, 95 kDa) in synaptosome (SN) compared to total homogenate (THF) fractions with respect to GAPDH (red, 37 kDa) loading control. 1 lane represents 1 mouse. **C)** Western blot validating levels of Aβ on freshly isolated synaptosomes treated with 50nM synthetic humanized oAβ 40-S26C compared to standard curve and GAPDH loading control. Aβ oligomers were mostly dimers (6.5 kDa) with some monomers (3kDa) in Aβ-

synaptosomes (Aβ-SN) with no immunoreactivity in control synaptosomes (Ctrl-SN). **D)** Immunocytochemistry showing increased levels of ePtdSer using PSVue 643 labelling (yellow) on oAβ-SN versus control Ctrl-SN immunostained with pre- and post-synaptic Synaptogamin 1/2 (magenta) and Homer1 (cyan) respectively. **E)** Western blot showing higher levels of Aβ dimer and monomer in synaptosomes prepared from the frontal cortex of AD (AD SN) patients and NDC (NDC SN) with respect to a standard curve with synthetic humanized Aβ 40-S26C and β-III-tubulin (55 kDa) loading control. 1 lane represents 1 preparation **(C)** or 1 patient **(E)**. Scale bar 500 nm **(A)** and 5 μm **(D)**.

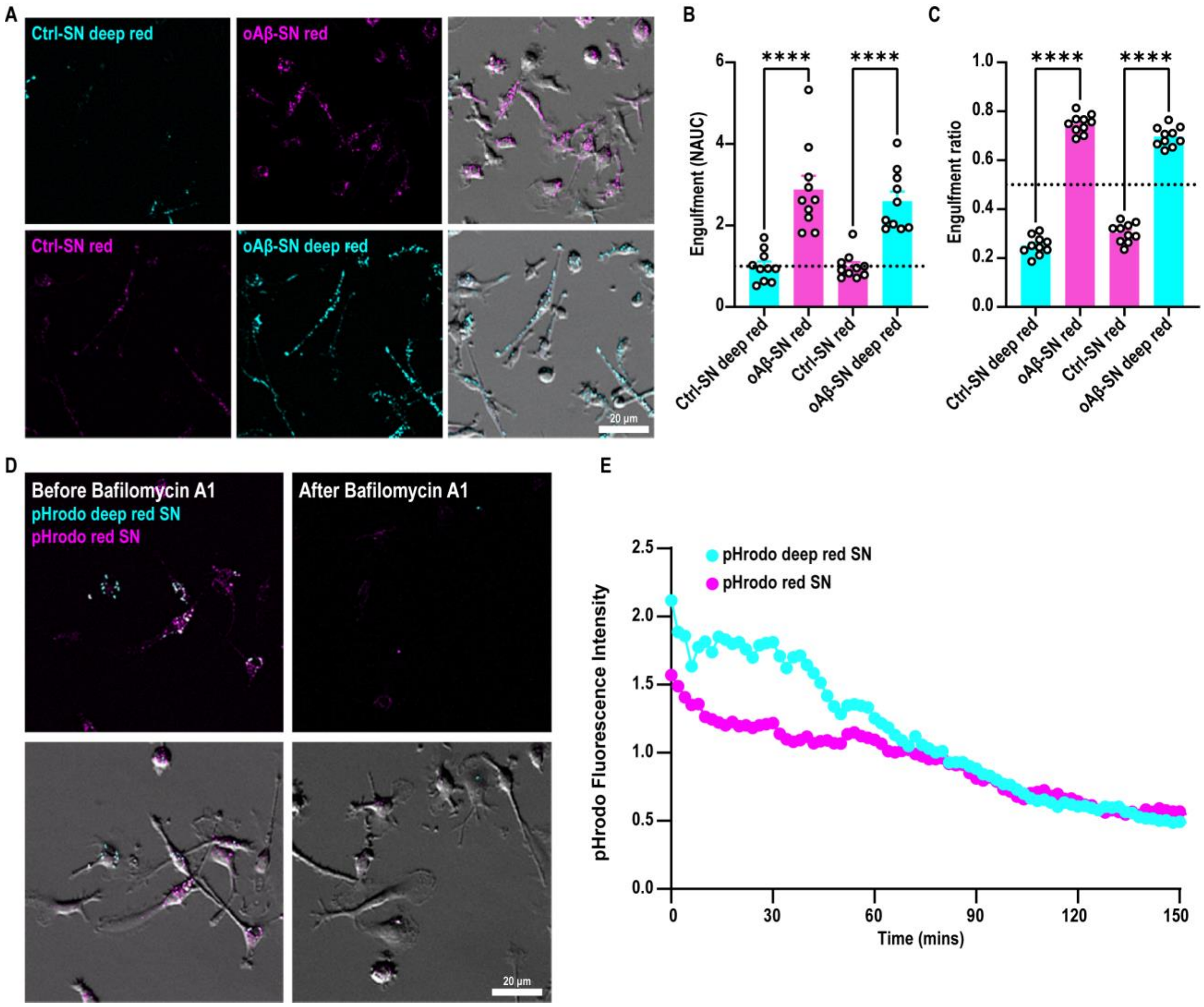


**Fig. S4. Microglia preferentially engulf oAβ-synaptosomes regardless of pHrodo dye.**
**A)** Primary microglia treated with simultaneously added oAβ-synaptosomes (oAβ-SN) and control synaptosomes (Ctrl-SN). Top panel: Ctrl-SN (cyan) and oAβ-SN (magenta) conjugated to deep red and red pHrodo respectively. Bottom panel: Ctrl-SN (magenta) and oAβ-SN (cyan) conjugated to red and deep red pHrodo respectively. **B)** pHrodo fluorescence intensity with time shown as area under curve (AUC) at 3 h normalized to respective control. AUC of oAβ-SN is higher than Ctrl-SN irrespective of pHrodo paradigm. **C)** Percent engulfed synaptosomes (SN) (oAβ-SN or Ctrl-SN/ total sum pHrodo fluorescence) at 3 h**. D)** Primary microglia treated simultaneously with synaptosomes conjugated to pHrodo red (red pHrodo SN, magenta) and deep red (deep red pHrodo SN, cyan) before and after 2 h treatment with bafilomycin A1 at the end of a microglia-synaptosome engulfment experiment. **E)** Bafilomycin, which prevents acidification of lysosomes, decreases pHrodo fluorescence intensity of both pHrodo red and deep red with time (2 min intervals, bafilomycin added at t=0). Data shown as mean ± SEM. 1 point represents 1 ROI. 6 ROIs, n=2 experiments per paradigm **(B-C)**. 2 ROIs **(E)**. Two-way ANOVA followed by Bonferroni post-hoc test, p-values shown as ****P<0.001. Scale bar 20 μm **(A, D).**

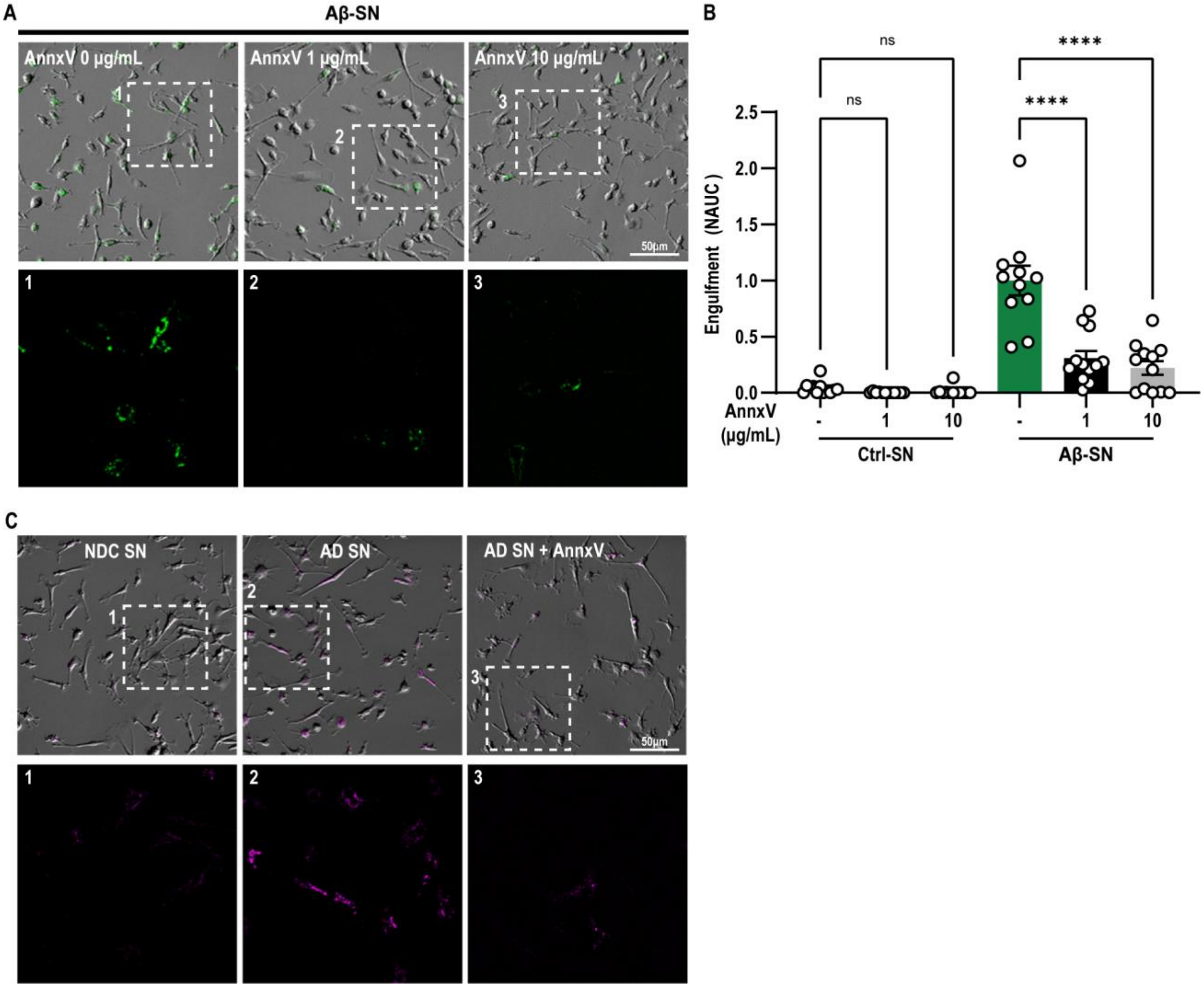


**Fig. S5. Annexin-V pre-treatment of AD synaptosomes decreases microglial engulfment.**
**A)** Primary microglia simultaneously treated with oAβ-synaptosomes (oAβ-SN) (green) conjugated to pHrodo red and control synaptosomes (Ctrl-SN) conjugated to pHrodo deep red pre-treated with either 0, 1 or 10 μg/mL of Annexin-V (AnnxV). **B)** pHrodo fluorescence intensity with time shown as area under curve (AUC) at 3 h. AnnxV treatment significantly decreases AUC of oAβ-SN but not Ctrl-SN. Data shown as normalized to non-treated oAβ-SN. **C)** Microglia treated with NDC (NDC SN), AD (AD SN) or AD synaptosomes pre-treated with 10 μg/mL AnnxV (AD SN + AnnxV) conjugated to pHrodo red (magenta). Data shown as mean ± SEM. 1 point represents 1 ROI (40 cells per ROI), 4 ROIs per experiment, n=3 experiments. Two-way ANOVA followed by Bonferroni post-hoc test, p-values shown as ****P < 0.001. Scale bars 50 μm **(A, C).**

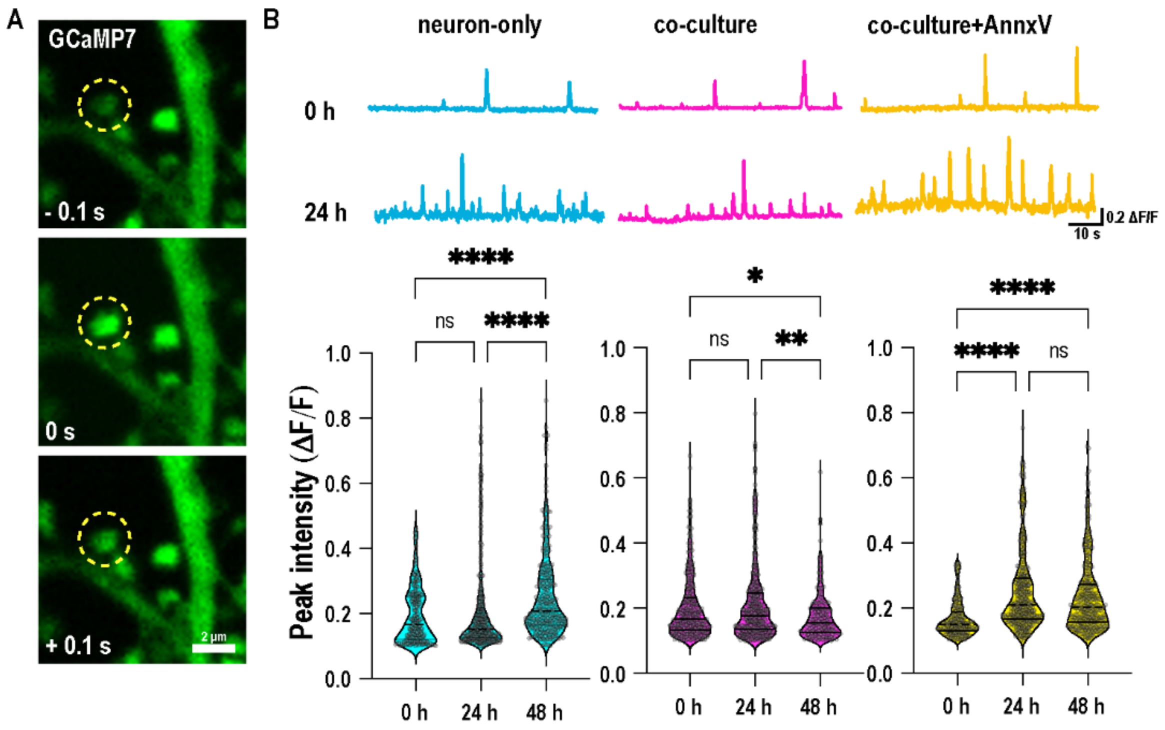


**Fig. S6. Co-culturing neurons with microglia resolves hyperactivity and ePtdSer.**

**A)** Representative image of transfected GCaMP7 signal on dendritic spines at 48 h post oAβ challenge. **B)** Representative trace (top panel) of GCaMP7 signal on dendritic spines at 0 h and 24 h. Box plots of normalized peak intensity ($\Delta F/F_0$) (bottom panel) of spontaneous GCaMP7 signal on dendritic spines of neuron-only (cyan), neuron-microglia co-culture (magenta) and neuron microglia co-culture treated with AnnxV (yellow) before (0 h) and after (24 h, 48 h) oAβ application. 1 point represents 1 ROI. 60-130 ROIs per experiment, n=3 experiments. Kruskal-Wallis test followed by Dunn's multiple comparisons test **(B)**. P-values shown as ns *P>0.05; **P<0.01; ****P<0.0001. Scale bar 2 µm **(A)**.

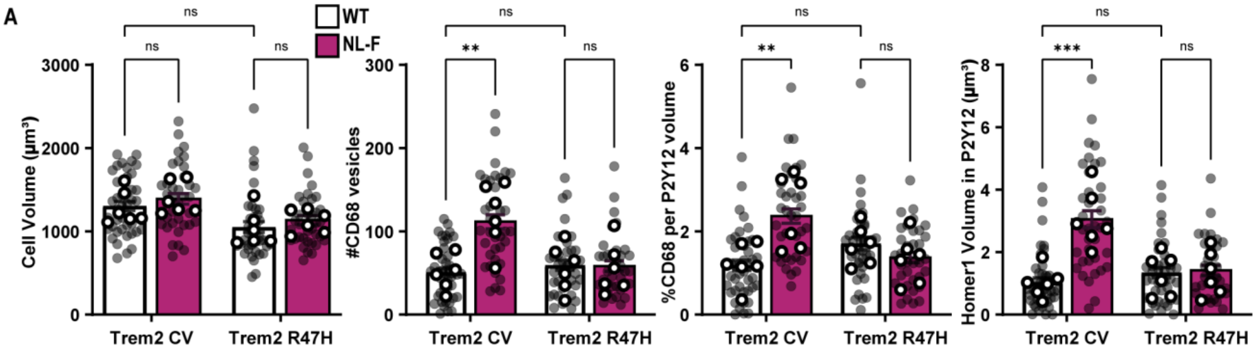


**Fig. S7. Microglia require functional TREM2 to engulf synapses in the NL-F AD model.**
**A)** Quantification of microglial P2Y12$^+$ cell volume, number of CD68$^+$ lysosomal vesicles per microglia, percentage of P2Y12$^+$ volume occupied by CD68$^+$ immunoreactive vesicles, and total volume of Homer1$^+$ material within P2Y12$^+$ microglia. All volumes are represented in μm$^3$. Data shown as mean ± SEM. Shaded points represent 1 microglial cell, open points represent animal average, 6-9 microglial cells per animal, n=6 animals. Two-way ANOVA followed by Bonferroni's post-hoc test, p-values shown as ns P>0.05; **P<0.01; ***P<0.001.

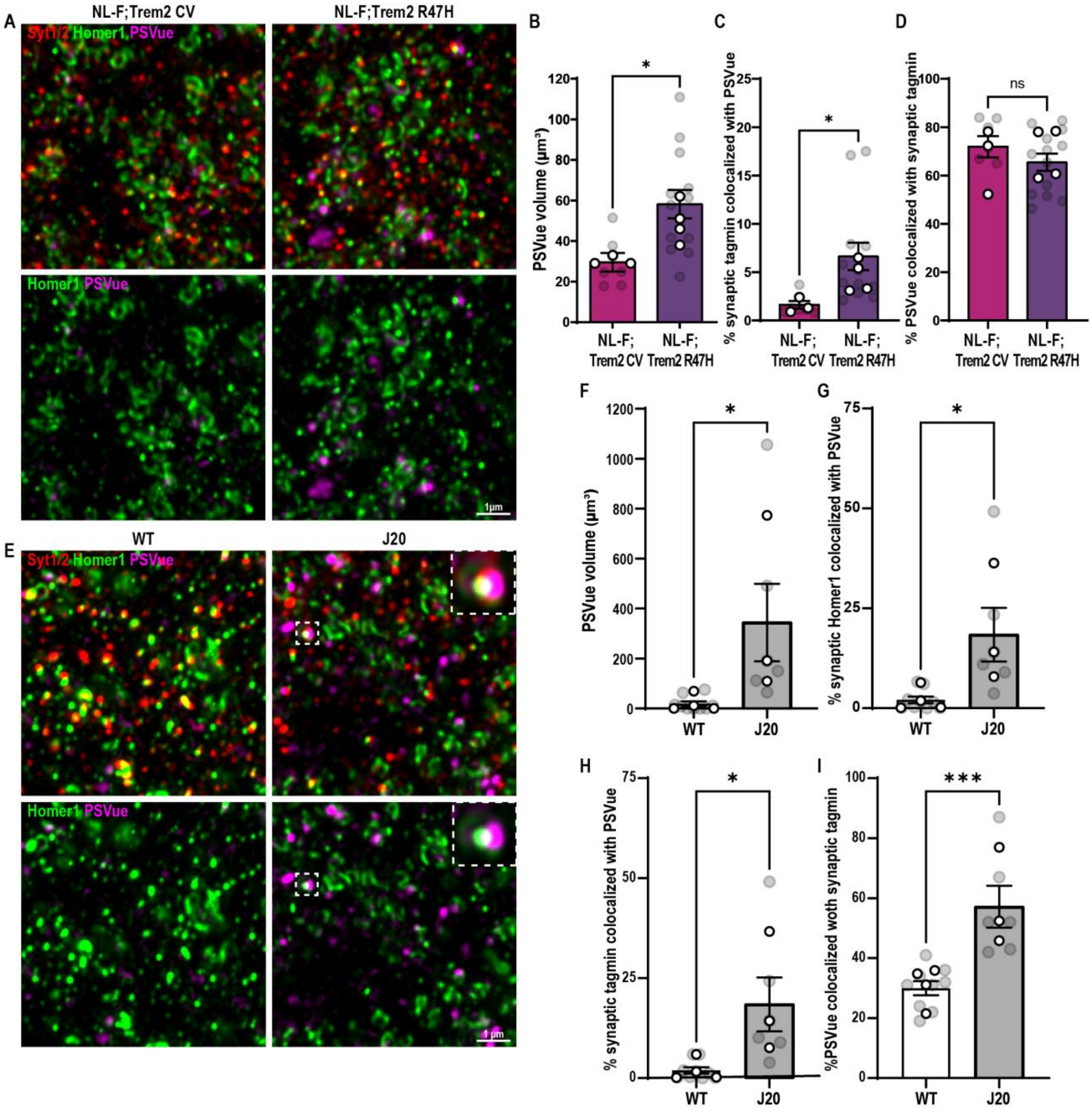


**Fig. S8. Trem2 loss-of-function exacerbates synaptic ePtdSer in the NL-F AD model.**

**A)** SRM images from the hippocampal CA1 dentate gyrus hilus of 6 mo NL-F KI and NL-F KI; Trem2 R47H KI mice immunostained for pre-synaptic Synaptotagmin 1/2 (Syt1/2, red) and post-synaptic Homer1 (green). PSVue 643 (magenta) is ICV injected. Upper panels show Syt1/2, Homer1 and PSVue. Lower panels show Homer1 and PSVue only. **B-C)** PSVue total volume per ROI **(B)** (7500 μm$^3$) and percentage of synaptic Homer1 puncta within 0.25 μm of PSVue **(C)** showing an increase in total volume and synaptic localized PSVue$^+$ respectively in NL-F KI; Trem2 R47H compared to NL-F KI. **D)** Percentage of PSVue volume within 0.25 μm of synaptic Synaptotamin 1/2 showing that approximately 80% of all PSVue immunostaining is localized to synapses. Note there is no difference between NL-F KI and NL-F KI; Trem2 R47H KI mice. **E)** SRM images from the hippocampal CA1 dentate gyrus hilus of 4 mo WT and J20 hAPP Tg mice immunostained for pre- Synaptotagmin 1/2 (Syt1/2, red) and post-synaptic Homer1 (green). PSVue 643 (magenta) is ICV injected. Left panels show Syt1/2, Homer1 and PSVue. Right panels show Homer1 and PSVue only. Insets show either triple (Syt1/2, Homer1 and PSVue) (left panel) or double colocalization (Homer1 and PSVue) (right panel). **F)** PSVue total volume per ROI (7500 $\mu m^3$) showing increased PSVue in J20 hAPP Tg mice compared to WT. **G-H)** Percentage of synaptic Homer1 **(G)** and synaptic Synaptotagmin 1/2 **(H)** puncta within 0.25 μm of PSVue showing an increase in $PSVue^+$ synapses in J20 hAPP Tg compared WT. **I)** Percentage of PSVue volume within 0.25 μm of synaptic Synaptotagmin 1/2 puncta. Data shown as mean ± SEM. Shaded points represent 1 ROI, open points represent animal average, 3 ROIs per animal, n=3-4 animals. Unpaired t-test, p-values shown as ns P>0.05; *P<0.05; ***P<0.001. Scale bar 1 μm.

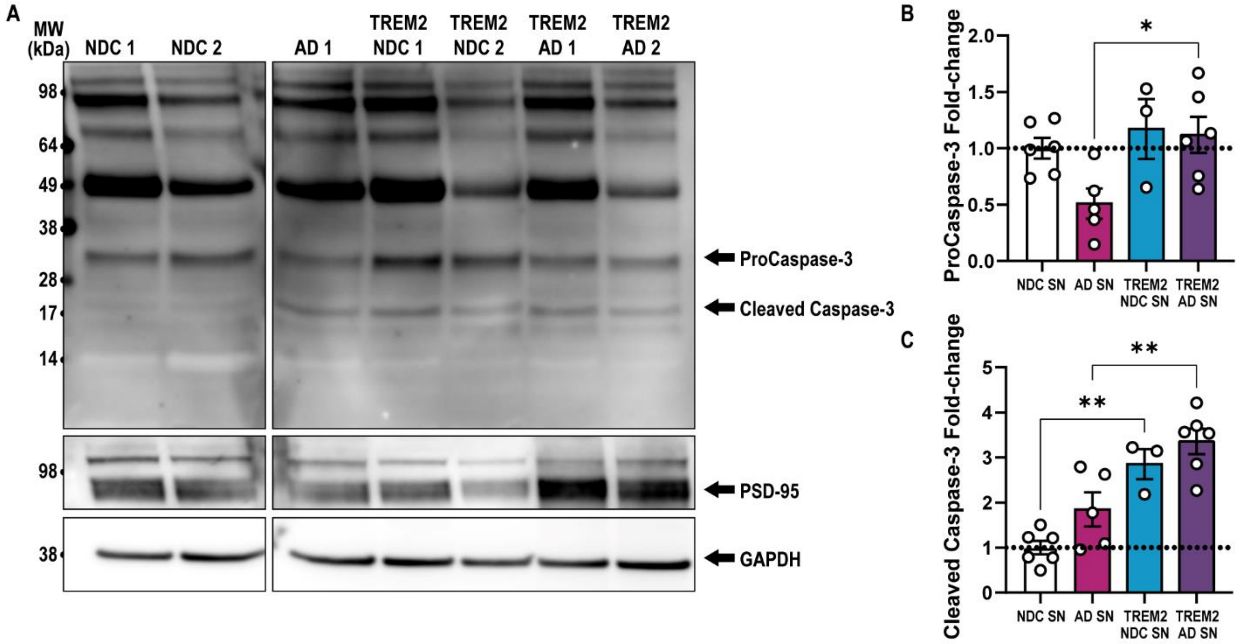


**Fig. S9. Increased caspase-3 activity on human AD synaptosomes.**

**A)** Western blots comparing levels of cleaved caspase-3 (17/19 kDa), procaspase-3 (35 kDa) and PSD-95 (95kDa) in human NDC, AD, NDC TREM2 and AD TREM2 synaptosomes (SN). 1 lane represents 1 patient. **B-C)** Western blot densitometry analysis showing levels of procasapse-3 **(B)** and cleaved caspase-3 **(C)** normalized to NDC. Data shown as mean ± SEM. 1 point represents 1 patient. One-way ANOVA followed by Bonferroni's post-hoc test, p-values shown as*P>0.05; **P<0.01.

### Table S1. Primer sequences

| Gene | Primer sequence 5’-3’ |
| --- | --- |
| *Actb* | Forward: CATTGCTGACAGGATGCAGAAGG, Reverse: TGCTGGAAGGTGGACAGTGAGG |
| *Cx3cr1* | Forward: GAGTATGACGATTCTGCTGAGG, Reverse: CAGACCGAACGTGAAGACGAG |
| *Gapdh* | Forward: CATCACTGCCACCCAGAAGACTG,Reverse: ATGCCAGTGAGCTTCCCGTTCAG |
| *Gfap* | Forward: CACCTACAGGAAATTGCTGGAGG, Reverse: CCACGATGTTCCTCTTGAGGTG |
| *Itgam* | Forward: ATGGACGCTGATGGCAATACC, Reverse: TCCCCATTCACGTCTCCCA |
| *Mag* | Forward: GGCCGAGGAGCAAGAATGG, Reverse: CATGCACTCTGCGATACGCT |
| *Map2* | Forward: ATGACAGGCAAGTCGGTGAAG, Reverse: CATCTCGGCCCTTTGGACTG |
| *Rpl32* | Forward: ATCAGGCACCAGTCAGACCGAT, Reverse: GTTGCTCCCATAACCGATGTTGG |
| *Tmem119* | Forward: CCTACTCTGTGTCACTCCCG, Reverse: CACGTACTGCCGGAAGAAATC |
| *Trem2* | Forward: CTGGAACCGTCACCATCACTC, Reverse: CGAAACTCGATGACTCCTCGG |

**Table S2. Human post-mortem frontal cortex samples.**

| ID | PMI | AO | AD | Gender | Clinical Diagnosis | Pathological Diagnosis | APOE | TREM2 | Braak | Thal | CERAD |
|---|---|---|---|---|---|---|---|---|---|---|---|
| 1 | 52h 05 | 58 | 68 | M | PCA | AD (PCA) | 34 | R62H | 6 | 5 | 3 |
| 2 | 35h 40 | 55 | 64 | M | CBD | AD | 34 | R47H | 6 | 5 | 3 |
| 3 | 51h 20 | 56 | 66 | F | AD | AD | 44 | R47H | 6 | 5 | 3 |
| 4 | 44h 00 | 59 | 75 | M | Picks | AD | 34 | R47H | 5 | 5 | 3 |
| 5 | 92h 20 | 58 | 62 | F | PCA | AD (PCA) | 34 | R62H | 6 | 5 | 2 |
| 6 | 81h 26 | 58 | 72 | F | AD | AD | 44 | R62H | 6 | 5 | 3 |
| 7 | 25h 30 | | 82 | M | Control | Cerebro-vascular disease | 33 | R47H | | | |
| 8 | 40h 10 | | 77 | M | Control | CAA | 22 | R47H | 0 | 0 | 0 |
| 9 | 76h 10 | | 71 | F | Control | Low level AD changes | 33 | R62H | 3 | 2 | 0 |
| 10 | 45h 05 | | 68 | F | Control | Control | 23 | - | 0 | 0 | 0 |
| 11 | 40h 20 | | 86 | F | Control | Control | 33 | - | 0 | 0 | 0 |
| 12 | 29h 30 | | 53 | F | Control | Cerebral infarct | 34 | - | 0 | 0 | 0 |
| 13 | 78h 50 | | 85 | M | Control | Control | 33 | - | 0 | 0 | 0 |
| 14 | 40h 10 | | 77 | M | Control | CAA | 22 | - | 0 | 0 | 0 |
| 15 | | | 86 | F | Control | | | - | 0 | 0 | 0 |
| 16 | | 60 | 71 | M | PPA | AD | 33 | - | 6 | 3 | 3 |
| 16 | | 84 | 91 | F | AD | AD | 34 | - | 4 | | 2 |
| 17 | | | 89 | M | Control | Control/ early AD | 33 | - | 2 | 3 | 1 |
| 18 | | | 87 | F | Control | Control/ early AD | 23 | - | 3 | 3 | 2 |
| 19 | | 88 | 88 | F | Control | AD | 33 | - | 4 | 5 | 2 |

**PMI:** Post-mortem interval

**AO:** Age of onset

**AD:** Age of death

**Table S3. Antibody information.**

| **Primary antibodies** | | | | |
|---|---|---|---|---|
| **Antibody Target** | **Catalogue No.** | **Company** | **Host** | **Dilution** |
| Homer1 | 160006 | Synaptic Systems | Chicken | 1/300-1/500 |
| Synaptotagmin 1/2 | 105002 | Synaptic Systems | Rabbit | 1/300 |
| 6E10 | 803001 | BioLegend | Mouse | 1/1000 |
| 4G8 | 800708 | BioLegend | Mouse | 1/1000 |
| GAPDH | ab181602 | Abcam | Rabbit | 1/20000 |
| Synaptophysin | ab8049 | Abcam | Mouse | 1/1000 |
| PDS-95 | MAB1596 | Merck | Mouse | 1/1000 |
| PSD-95 | 124014 | Synaptic Systems | Guinea pig | 1/1000 |
| Cleaved caspase-3 | 9661S | Cell Signalling | Rabbit | 1/1000 |
| Caspase-3 | 9662 | Cell Signalling | Rabbit | 1/1000 |
| IBA1 | 019-19741 | Wako Chemicals | Rabbit | 1/500 |
| P2Y12 | AS-55043A | Anaspec | Rabbit | 1/500 |
| CD68 | MCA-1957 | Serotec | Rat | 1/500 |
| TMEM119 | ab209064 | Abcam | Rabbit | 1/1000 |
| **Secondary antibodies** | | | | |
| **Fluorophore tag** | **Catalogue No.** | **Company** | **Host** | **Dilution** |
| Anti-Chicken 488 | A11039 | ThermoFisher | Goat | 1/500 |
| Anti- Rabbit 594 | A11037 | ThermoFisher | Goat | 1/500 |
| Anti-Rabbit 546 | A11035 | ThermoFisher | Goat | 1/500 |
| Anti-Rabbit 647 | A27040 | ThermoFisher | Goat | 1/1000 |
| Anti-Rat 647 | A21247 | ThermoFisher | Goat | 1/500 |
| Anti- Mouse HRP | ab205719 | Abcam | Goat | 1/5000 |
| Anti- Rabbit HRP | ab205718 | Abcam | Goat | 1/10000 |
| Anti-Mouse 800 | A32789 | ThermoFisher | Goat | 1/1000 |
| Anti-Rabbit 680 | A32734 | ThermoFisher | Goat | 1/1000 |

**Movie S1. Microglia contact ePtdSer⁺ dendritic spines.**

Time-lapse video of microglia (yellow, labelled with IB4-647, 3D rendered) co-cultured with Homer1-eGFP hippocampal neurons (green) treated with 50nM oAβ. Microglia contact PSVue$^+$ (magenta) Homer-1eGFP dendritic spines. Scale bar 5 μm.

**Movie S2. Microglia internalize ePtdSer$^+$ dendritic spines.**

Time-lapse video of microglia (blue, labelled with IB4-647, 3D rendered) co-cultured with Homer1-eGFP hippocampal neurons (green) treated with 50nM oAβ. Microglia internalize PSVue$^+$ (magenta) Homer-1eGFP dendritic spines. Scale bar 5 μm.

**Movie S3. Microglia preferentially engulf Aβ-synaptosomes over control.**

Time-lapse video of primary microglia preferentially engulfing Aβ-synaptosomes in pHrodo red (magenta) over control synaptosomes in pHrodo deep red (cyan) over 10 h (2-5 min intervals). Scale bar 50 μm.

**Movie S4. Spontaneous calcium transients in GCaMP7 transfected hippocampal neurons.**

Normalized fluorescence of dendritic spines of neurons transfected with GCaMP7. A non-linear scale was used to show the fluorescence intensity, ranged from low (dark blue) to high values (white). Scale bar 5 μm.

**Movie S5. No overt changes in motility and morphology in Trem2 R47 KI microglia.**

Time-lapse video of primary microglia prepared from Trem2 R47H KI mice showing no overt changes microglial morphology and motility. Scale bar 50 μm.
